# Trymethylamine-N-oxide, a gut-derived metabolite, induces myofibroblastic activation of valvular interstitial cells through endoplasmic reticulum stress

**DOI:** 10.1101/2025.02.06.636980

**Authors:** Samanvitha Sudi, Sai Drishya Suresh, Tanmayee Kolli, Ana Maria Porras

## Abstract

Calcific aortic valve disease currently lacks effective treatments beyond surgical valve replacement, due to an incomplete understanding of its pathogenesis. Emerging evidence suggests that the gut microbiome influences cardiovascular health through the production of metabolites derived from dietary components. Among them, trimethylamine-N-oxide (TMAO) has been identified as a potential causal factor for several cardiovascular conditions. However, its role in the development of aortic valve disease remains poorly understood. This study sought to investigate the impact of TMAO on valvular interstitial cells (VICs), the most abundant cell type in the aortic valve. Here, we demonstrate that TMAO activates VICs towards a myofibroblastic profibrotic phenotype. Using an *in vitro* protocol to generate quiescent VICs, we found that TMAO induces the upregulation of myofibroblastic markers in a sex-independent manner. These quiescent VICs were more sensitive to TMAO than conventionally cultured VICs. Treatment with TMAO also elevated extracellular matrix production and oxidative stress, phenotypic hallmarks of an activated profibrotic state. Finally, inhibition of the endoplasmic reticulum stress kinase prior to TMAO treatment blocked all effects of this metabolite. These findings suggest that TMAO contributes to the early stages of valve disease by promoting VIC activation through endoplasmic reticulum stress mechanisms. Understanding the role of TMAO and other gut-derived metabolites in the pathogenesis of valve disease could inform the development of novel preventive or therapeutic strategies to modify or delay disease progression. Furthermore, these insights underscore the importance of host-microbiome interactions and highlight the potential for targeted dietary interventions to mitigate cardiovascular disease risk.

## INTRODUCTION

Despite the quadrupling of the global prevalence of calcific aortic valve disease (CAVD) over the last three decades [1], surgical valve replacement remains the sole treatment option [2,3]. The development of novel pharmacological treatments depends on the identification of the cellular and molecular mechanisms involved in the pathogenesis of valvular disease. Accumulating evidence indicates the gut microbiome influences cardiovascular health, partly through the production of metabolites derived from dietary components [4–6]. While these microbial metabolites have been identified as key mediators in the pathogenesis of various cardiovascular conditions [7,8], their potential roles in the progression of CAVD remain incompletely understood.

Trimethylamine N-oxide (TMAO) has emerged as a gut-derived metabolite of particular interest due to its association with multiple cardiovascular diseases [9–12]. TMAO is produced through a two-step process. First, gut bacteria metabolize dietary micronutrients present in high-fat foods, such as choline and carnitine, generating trimethylamine [10,13,14]. This intermediate is then transported to the liver, where hepatic flavin monooxygenases oxidize it to TMAO, subsequently entering the systemic circulation [15–17]. In clinical studies, high circulating TMAO levels have been associated with increased risk of major adverse cardiovascular events [18], coronary heart disease [19], cardiometabolic disease [20], hypertension [21], and atherosclerosis [22]. Furthermore, animal and *in vitro* studies implicate TMAO as a causal factor in the development of cardiovascular disease through mechanisms involving endothelial dysfunction [23–25], inflammation [26], foam cell formation [27], and fibrosis [28].

Increasing evidence suggests that TMAO may be involved in the progression of aortic valve sclerosis and calcification [29–32]. In a retrospective clinical study, patients with severe aortic stenosis had higher serum levels of TMAO compared to control subjects, even after adjusting for baseline characteristics [32]. Increased TMAO was also associated with poor adverse outcomes after transcatheter aortic valve replacement [32]. A similar cohort study found elevated TMAO to be a predictor of CAVD [31]. Moreover, in recent *in vivo* mouse studies, supplementation with dietary choline led to elevated TMAO levels, thickened aortic valves, and valvular fibrosis [30,31]. Treatment with 3,3-dimethyl-1-butanol, an inhibitor of TMA formation, prevented these effects [30,31]. Collectively, these clinical and animal data suggest a causal connection between TMAO and CAVD, with further research necessary to precisely define the contributions of this gut metabolite to the initiation of valve disease.

One of the early events in the development of CAVD is the activation of valvular interstitial cells (VICs), which maintain the structural integrity and function of the valve [33]. In a healthy state, these fibroblast-like cells remain quiescent [34]. However, during the progression of CAVD, qVICs undergo phenotypic changes, transitioning first into activated myofibroblasts and later into osteoblastic-like cells [35]. This myofibroblastic transformation marks a crucial initial step in the cascade of pathological events during the early stages of valve disease [36,37]. Others have shown that TMAO can modulate VIC phenotype *in vitro* towards fibrotic [30] and osteoblastic [31] behavior. Additionally, Li *et al.* demonstrated that TMAO can induce cardiac fibroblast activation marked by increases in proliferation, migration, and collagen deposition [28]. Hence, we hypothesized that TMAO may contribute to the early stages of valve disease by activating quiescent VICs (qVICs).

Testing this hypothesis through conventional cell culture techniques is challenging because VICs spontaneously activate when cultured on traditional tissue culture plastic [36,37]. As a result, these *in vitro* culture approaches that predominantly exhibit the activated (aVIC) phenotype may not reliably represent the process of VIC activation or the characteristics of a healthy valve. To address this issue, we have previously developed a protocol to generate qVICs in 2D *in vitro* culture [38]. Using this protocol, we sought to assess the impact of TMAO on healthy quiescent VICs. Here, we demonstrate that TMAO triggers qVIC activation towards a profibrotic myofibroblastic phenotype independent of cellular sex. Furthermore, we show that qVICs are more sensitive than aVICs to this gut metabolite. Finally, we determined that TMAO induces VIC activation by promoting endoplasmic reticulum (ER) stress mediated by the endoplasmic reticulum stress kinase (PERK) pathway.

## MATERIALS AND METHODS

### VIC Isolation, qVIC Generation, and TMAO Treatment

Aortic valve leaflets were harvested from male and female (6 to 9 months) porcine hearts (Animal Technologies, Tyler, Texas). Male and female cells were cultured separately throughout the expansion and all experiments. Aortic valve leaflets were excised and thoroughly washed in a heart wash solution containing deionized water, M199 powder, 2% penicillin/streptomycin, and 1% L-glutamine. The leaflets were then incubated at 37°C for 30 minutes in a collagenase II solution (Worthington Biochemical Corporation). Following incubation, the valves were vortexed for 30 seconds to dislodge the valvular endothelial cells. After aspirating the solution, the remaining undigested tissue was removed, and placed in fresh collagenase II solution at 37°C. After a 2 hour incubation, the leaflets were vortexed for 2 minutes to dislodge the VICs, which were then passed through a 100μm cell filter for filtration. The VIC suspension was centrifuged at 500 RCF, and the resulting cell pellet was resuspended, and plated on tissue culture flasks in low glucose DMEM supplemented with 10% FBS and 1% penicillin/streptomycin.

Upon reaching confluency, qVICs were generated following our previously established protocol [38]. Briefly, VICs were passaged onto collagen-coated (2 μg/cm^2^; Human Collagen Solution, Type III, Advanced Biomatrix) tissue culture polystyrene (TCPS) and cultured in low glucose DMEM supplemented with 2% FBS, 1% penicillin/streptomycin, 5.25 μg/mL insulin and 10 ng/mL fibroblast growth factor (FGF; Peprotech) for 10 days. To generate aVICs, the cells were cultured on uncoated TCPS in low glucose DMEM supplemented with 10% FBS.

For all experiments, qVICs and aVICs were seeded at a density of 10,000 cells/cm^2^ and cultured in DMEM supplemented with 2% FBS. Twenty-four hours after seeding, VICs were treated with TMAO (Thermo Scientific) at concentrations ranging from 25 µM to 150 µM. VICs were re-fed and treated again with TMAO every 48 hours. Untreated VICs served as the negative control, while treatment with transforming growth factor beta (TGF-β_1_; 0.5 ng/mL for qVICs and 5 ng/mL for aVICs) served as the positive control. VIC phenotype was analyzed after 3 and 5 days of treatment.

### Immunocytochemistry for Myofibroblastic Markers

The expression of proteins associated with the myofibroblastic phenotype was analyzed via immunocytochemistry. VICs were washed twice with phosphate-buffered saline (PBS) and fixed with 10% formalin for 10 minutes. After formalin removal, the cells were washed with PBS and permeabilized with 0.5% Triton X for 5 minutes before blocking with 3% bovine serum albumin (BSA) for 1 hour. Following blocking, the VICs were incubated with anti-αSMA primary antibody (monoclonal, clone 1A4; Sigma Aldrich) diluted 1:500 in 1% BSA or anti-smooth muscle protein 22-α (SM22; Abcam) diluted 1:500 in for 90 minutes at room temperature. The primary antibody was then removed, and the cells were washed twice with PBS. Next, the samples were incubated with an Alexa Fluor 555 goat anti-mouse secondary antibody (Invitrogen) in 1% BSA (1:1000) for 60 minutes at room temperature. After three washes with PBS, DAPI staining was applied for 5 minutes to label the nuclei, followed by an additional three PBS washes. Fluorescent images were captured using a BZ-800 Keyence microscope. Integrated fluorescent intensity per field of view was quantified with BZ analysis software and normalized to the corresponding DAPI-stained nuclei count.

### Gene Expression Analysis with qRT-PCR

To evaluate gene expression levels, quantitative reverse transcription–polymerase chain reaction (qRT-PCR) was employed. VIC RNA was extracted using the RNEasy Mini Kit (Qiagen) and converted into cDNA using the High-Capacity cDNA Reverse Transcription Kit (Applied Biosystems). qRT-PCR was then performed with TaqMan Gene Expression Assays (Applied Biosystems) for myofibroblastic markers (αSMA [ACTA2] and Transgelin [SM22]), extracellular matrix proteins (fibronectin and collagen type I [COL1A1]), and osteoblastic markers (alkaline phosphatase [ALP] and bone morphogenetic protein [BMP]). The comparative CT (ΔΔCT) method was applied, with gene expression for each experimental condition normalized to the endogenous control GAPDH and then compared to the untreated control condition. In comparing male and female VICs, the gene expression levels were compared to those of the untreateted female control.

### Quantification of Proliferation and Apoptosis

Cell proliferation was analyzed using the Click-iT EdU Alexa Fluor 488 Cell Proliferation Imaging Kit (Invitrogen). VICs were treated with EdU for 8 hours, fixed, permeabilized, and labeled with Alexa Fluor 488 per the kit’s instructions. Cell nuclei were stained with DAPI, and fluorescent images were acquired using a Keyence BZ-800 microscope. Positive EdU-labeled cells were quantified using the Keyence BZ analyzer software and normalized to the total number of cells labeled with DAPI to obtain the percentage of proliferating cells.

Apoptosis was measured with the SenosoLyte Homogeneous AFC Caspase 3/7 Assay Kit (AnaSpec). In brief, VICs were incubated with Caspase-3/7 substrate for 60 minutes at 37°C. End-point fluorescence intensity readings were acquired with a SpectraMax microplate reader (Molecular Devices) at an excitation wavelength at 500nm and normalized to the average DAPI cell count per well for the corresponding condition.

### Quantifying collagen and fibronectin secretion and deposition through ELISAs

Fibronectin deposition was analyzed semi-quantitatively using an *in situ* ELISA. After 5 days of TMAO treatment, VICs were fixed with 10% formalin for 10 minutes followed by treatment with hydrogen peroxide (0.3% v/v in methanol) for 1 hour to quench endogenous peroxidase activity. Subsequently, samples were blocked with 3% BSA overnight and stained on the next day with a mouse anti-fibronectin antibody (1:500; monoclonal IgG1; Santacruz) in 1% BSA for 2 hours at room temperature. After removing the primary antibody, the cells were washed 3 times with PBS. The primary antibody was then labeled with horseradish peroxidase–linked goat secondary antibody (Goat anti-Mouse IgG Secondary Antibody; Thermo Fisher Scientific) diluted 1:1000 in 1% BSA for 40 minutes. After additional washing with PBS, 1-Step^TM^ Turbo TMB-ELISA substrate solution (Thermo Scientific) was added for 5 minutes, and the reaction was stopped with 2N sulfuric acid. Absorbance at the 450 nm wavelength was measured using a SpectraMax microplate reader. To account for background signal, 2 wells per condition were incubated with no primary antibody. The absorbance from these background controls was subtracted from the total absorbance for each condition. Finally, the absorbance values were normalized based on the previously calculated average DAPI cell count for each condition.

We also quantified fibronectin and collagen secretion in the culture supernatant via DuoSet ELISAs (R&D Systems). VIC culture supernatant was collected 3 days post-treatment, and the ELISAs were performed according to the guidelines provided by the manufacturer. Absorbance was measured (λ = 450 nm) using a SpectraMax plate reader. The calculated concentration of fibronectin or collagen in each condition was normalized to the average DAPI cell count for that condition.

### Reactive Oxygen Species Assessment

VIC generation of reactive oxygen species was quantified using the ROS-Glo™ H₂O₂ Assay kit (Promega). Briefly, the cells were incubated with H₂O₂ substrate for 6 hours at 37°C. Later, a luciferin-based detection solution was added and incubated for 20 minutes. Luminescence was measured using a SpectraMax plate reader and normalized to average DAPI cell counts. ROS levels were also assessed using the Cellular ROS Assay Kit (Abcam) VICs were incubated with the cell-permeant reagent, 2’,7’-dichlorofluorescin diacetate, for 45 minutes. This substrate fluoresces in the presence of reactive oxygen species. Fluorescence images were captured with a Keyence BZ microscope. Integrated fluorescent intensity per field of view was quantified with BZ analysis software and normalized to the corresponding DAPI-stained nuclei count.

### Evaluation of Endoplasmic Reticulum Stress

Endoplasmic Reticulum (ER) stress was analyzed using the ER-ID® Red assay kit (Enzo Life Sciences). Briefly, cells were washed twice with PBS and fixed in 10% formalin for 10 minutes. Following fixation, the cells were washed again with PBS and permeabilized with 0.5% Triton X-100 for 5 minutes. The cells were then incubated with ER-ID® Red and Hoechst 33342 nuclear stain for 30 minutes at room temperature. After incubation, the cells were washed three times with the provided assay buffer. Fluorescent images were captured using a BZ-800 Keyence microscope. The integrated fluorescent intensity per field of view was quantified using BZ analysis software and normalized to the Hoechst-stained nuclei count.

### Inhibition of the PERK Pathway and TGF-β_1_ Receptor

qVICs were seeded at a density of 10,000 cells/cm^2^ in low glucose DMEM containing 2% FBS. After 24 hours, the cells were treated with GSK2656157 (1μM; Santa Cruz), an inhibitor of the endoplasmic reticulum stress kinase (PERK) pathway, or SB431542 (Selleckchem), an inhibitor of the TGF-β type I receptor, for 1 hour at 37°C. Following this initial treatment, the cells were treated with 150 µM TMAO for 3 days. At the end of the experiment, cell proliferation, the expression of myofibroblastic markers, ROS production, and ECM deposition were assessed.

### Statistical Analysis

Each experiment was performed using VICs isolated and pooled from 3 to 4 individual porcine hearts separated by sex with n=4-6 technical replicates per condition. One-way ANOVA followed by Tukey’s multiple comparisons test was used for all analyses, except where comparisons involved the effects of the PERK inhibitor or sex-based responses to treatment. In those cases, a two-way ANOVA was applied. All the statistical analyses were conducted using GraphPad Prism software, and the data are presented as mean ± standard deviation.

## RESULTS

### TMAO activates quiescent valvular interstitial cells

First, we generated quiescent VICs (qVICs) to test the hypothesis that TMAO triggers VIC activation. Primary female porcine VICs were cultured on collagen-coated plates (2 µg/cm^2^) in media supplemented with 2% fetal bovine serum (FBS), insulin (5.25 μg/mL), and fibroblast growth factor (10 ng/mL) for 10 days as previously reported [38]. The generated qVICs were then treated with varying concentrations of TMAO (25 to 150 µM) for 3 days (Fig 1A). For these experiments, two positive controls were included – VICs treated with TGF-β_1_ (1 ng/mL), a known stimulus of myofibroblastic differentiation [39], and activated VICs (aVICs) generated through standard culture on uncoated TCPS in media supplemented with 10% FBS.

**Figure 1.**
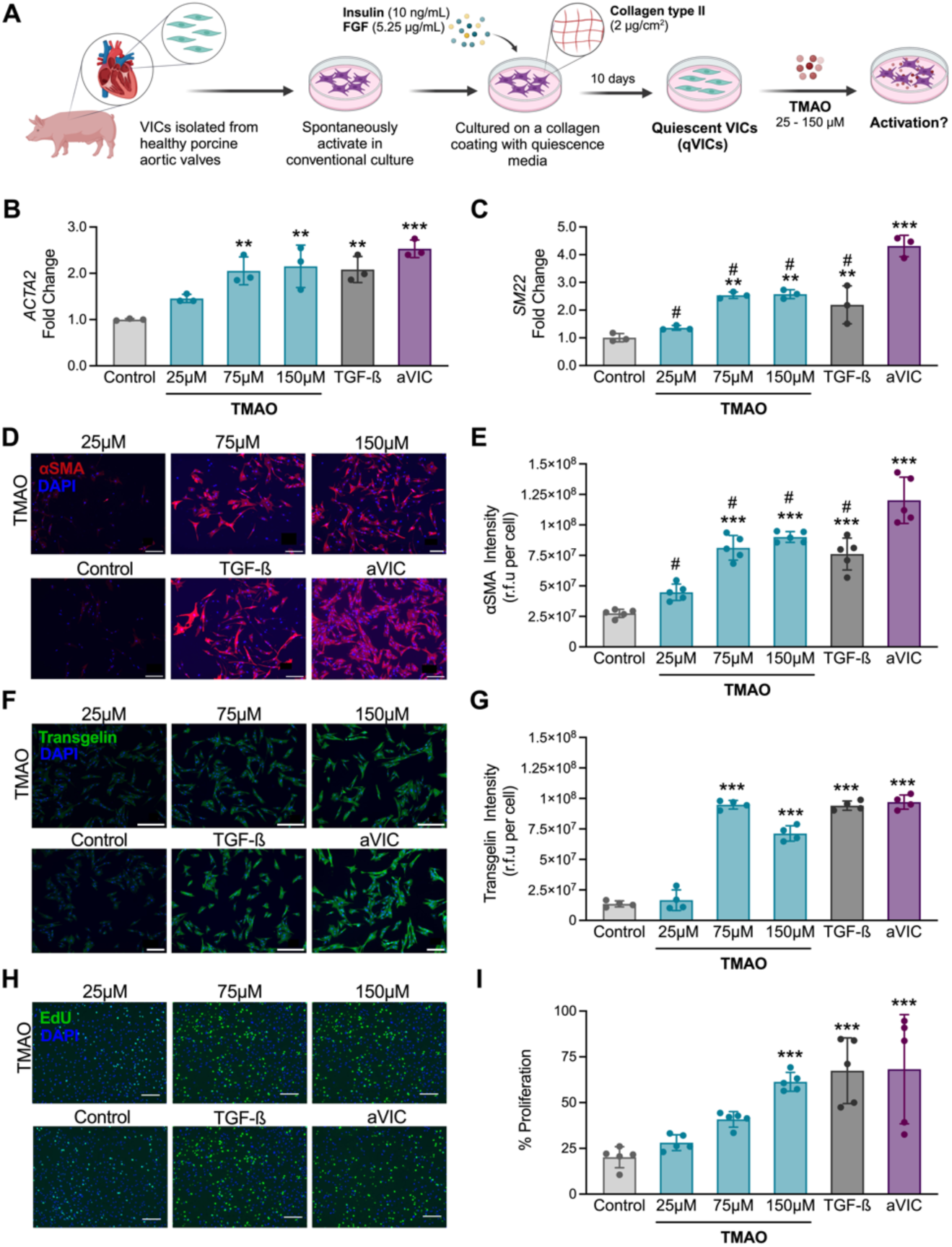
TMAO triggers qVIC activation. (A) Schematic illustration of the experimental design. Primary VICs were isolated from porcine aortic valves and cultured to generate quiescent VICs (qVICs). qVICs were treated with TMAO for 3 days. (B-C) Quantification of myofibroblastic gene expression levels for (B) *ACTA2* and (C) *SM22* via qRT-PCR. (D) Representative images of immunocytochemistry staining for αSMA (red). Cell nuclei are stained in blue. (E) Quantification of αSMA staining intensity in (D). (F) Representative images of immunocytochemistry staining for transgelin (green). Cell nuclei are stained in blue. (G) Quantification of transgelin staining intensity in (E). (H) EdU staining to identify proliferating cells after 8 hours of incubation with EdU. (I) Quantification of the percentage of EdU-positive proliferating cells in (H). Scale bars represent 200 µm. n = 3-5 replicates per condition. One-way ANOVA followed by Tukey’s multiple comparisons test. \*\**p*<0.005, \*\*\**p*<0.001 compared with the control. #*p*<0.01 compared with aVICs.

Treatment with TMAO at concentrations equal to or above 75 µM led to significant upregulation of *ACTA2* (Fig 1B) and *SM22* (Fig 1C), known gene markers for the activated myofibroblastic phenotype [40]. Significant increases in the expression levels of the corresponding proteins, αSMA (Fig 1D-E) and transgelin (Fig 1F-G) were also observed via immunocytochemistry, confirming the phenotypic shift toward a myofibroblastic state after TMAO treatment. Additionally, TMAO-treated qVICs exhibited α-SMA and transgelin expression levels comparable to those observed after treatment with TGF-β_1_ (Fig 1A-G). qVICs treated with 150 µM TMAO also displayed increased proliferation rates, equivalent to those of both activation controls (Fig 2H). No significant differences in apoptosis were observed across any treatment groups (S1 Fig). The increase in both the expression of myofibroblastic markers and proliferation indicates that TMAO indeed promotes the transition of qVICs to an activated state.

**Figure 2.**
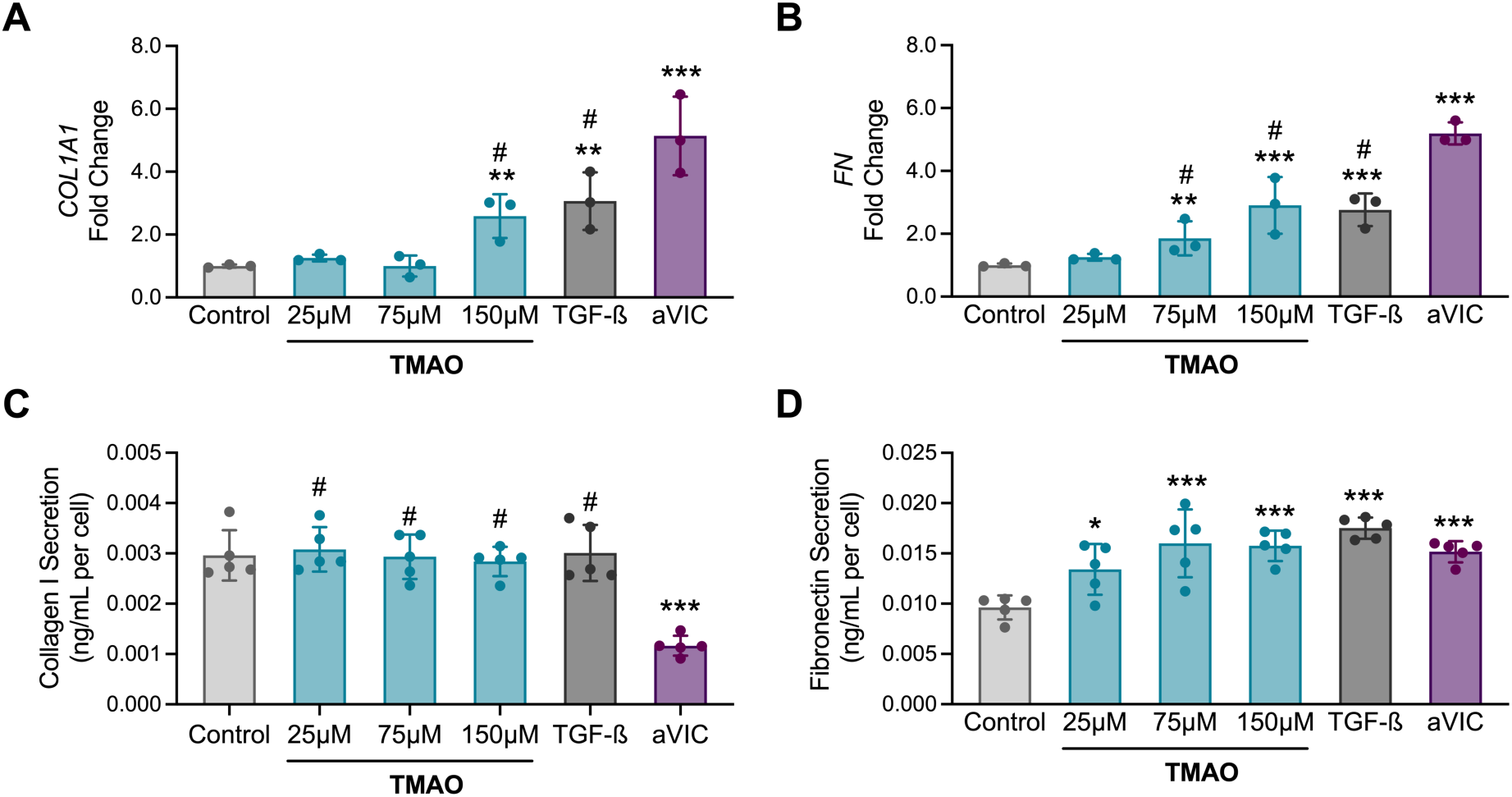
TMAO increases extracellular matrix production by qVICs. (A-B) Gene expression analysis of (A) *COL1A1* and (B) *FN* after 3 days of TMAO treatment via qRT-PCR. (C-D) Quantification of (C) collagen and (D) fibronectin secretion via sandwich ELISAs. n = 3-5 replicates per condition. \*\**p*<0.005, \*\*\**p*<0.001 compared with the control, #*p*<0.05 compared to aVICs.

### Treatment with TMAO induces extracellular matrix (ECM) production

Because the activation of qVICs into myofibroblastic-like cells is often associated with increased ECM deposition [35], we also evaluated the impact of TMAO treatment on the production of two ECM proteins: collagen I and fibronectin. After 3 days of treatment, TMAO significantly upregulated the gene expression of *COL1A1* (Fig 2A) and *FN* (Fig 2B) in qVICs, reaching levels comparable to those observed after treatment with TGF-β_1_. The upregulation of *COL1A1* was observed only at the highest concentration of 150 µM, while *FN* was upregulated at 75 µM and higher concentrations. To account for potential post-transcriptional modifications and protease degradation [41], we also assessed the production of these components at the protein level. No statistically significant differences were observed in collagen secretion (Fig 2C) or deposition (S2A Fig) following TMAO treatment. In contrast, TMAO treatment significantly increased fibronectin secretion at all concentrations (Fig 2D) and deposition at concentrations ^3^≥ 75 µM (S2B Fig).

We next sought to examine the effects of TMAO on conventionally cultured activated VICs (Fig 3A). For these experiments, the control consisted of untreated aVICs. aVICs treated with TGF-β_1_ (10 ng/mL) and untreated qVICs served as additional positive and negative controls, respectively. Initially, female aVICs were treated with the same concentrations used for qVICs (25-150 µM). However, TMAO had no effect on α-SMA expression (S3A-B Fig) or proliferation (S3C Fig) at these concentrations. Thus, we proceeded to treat the aVICs with higher TMAO concentrations (300 µM and 600 µM; Fig 3A). After 3 days of treatment, we observed a significant upregulation of the *ACTA2* gene at 600 µM (Fig 3B), while the *SM22* gene was upregulated at concentrations of 300 µM and 600 µM (Fig 3C). However, this increase in myofibroblastic marker expression was not reflected at the protein level for either α-SMA (Fig 3D-E) or transgelin (S4A-B Fig). At these higher concentrations, TMAO led to a small decrease in cell proliferation for all concentrations (S4C-D Fig).

**Figure 3.**
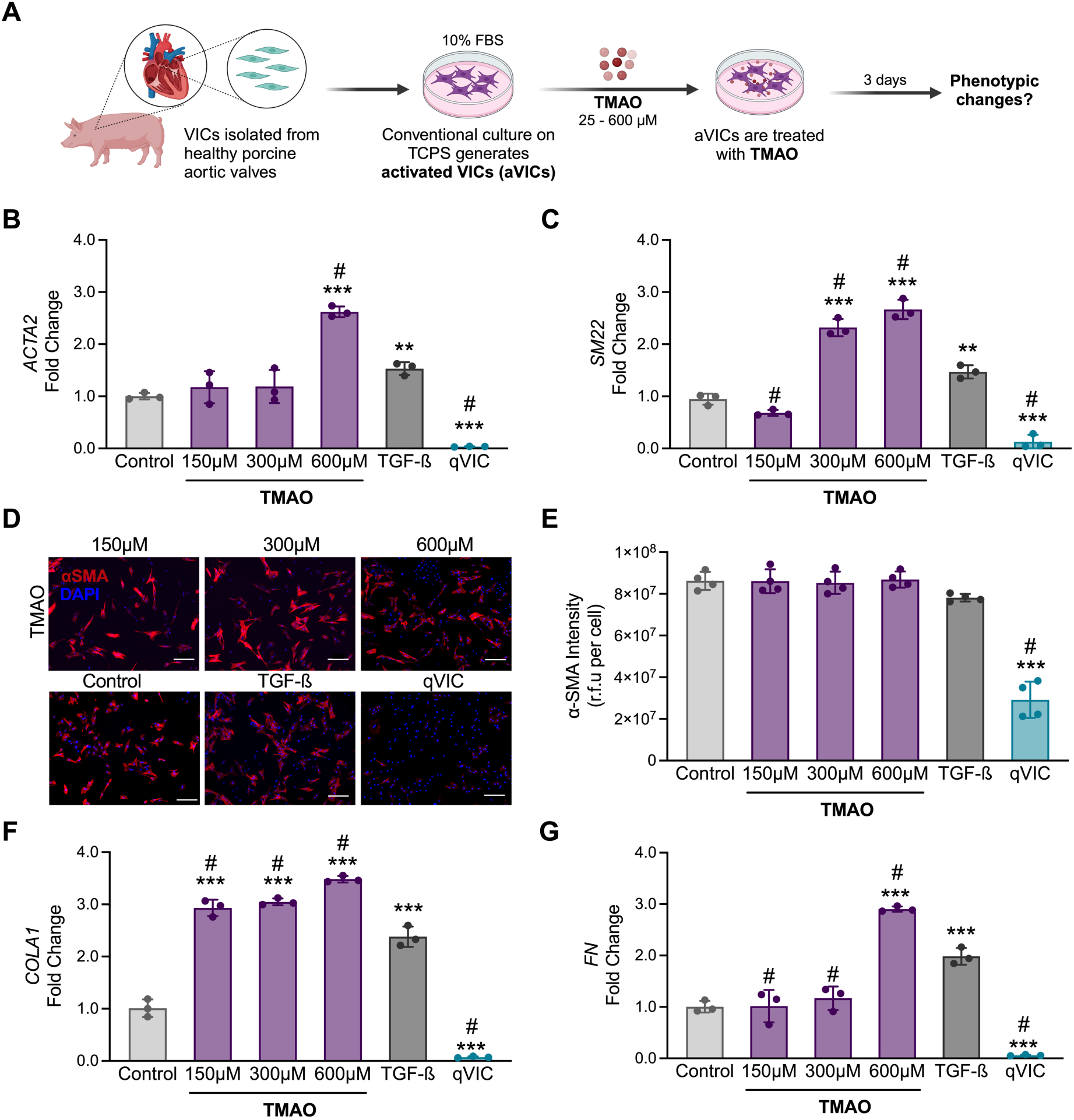
Effects of TMAO on conventionally cultured aVICs. (A) Schematic illustration describing the generation of activated VICs (aVICs) from primary porcine cells through conventional culture. aVIC were treated with TMAO for 3 days. (B-C) Gene expression of (B) *ACT2* (C) *SM22* via qRT-PCR. (D) Immunocytochemistry staining for αSMA. The scale bar represents 200 µm. Cell nuclei are stained blue. (E) Quantification of αSMA staining intensity in (D). (F-G) Analysis of (F) *COL1A1* and (G) *FN* gene expression via qRT-PCR*. n* = 3-5 replicates per condition. One-way ANOVA followed by Tukey’s multiple comparisons test. \*\**p*<0.005, \*\*\**p*<0.001 compared with the control. #*p*<0.01 compared to treatment with TGF-ß_1_.

Finally, we evaluated the production of ECM proteins by aVICs treated with TMAO. Treatment with high concentrations of TMAO resulted in statistically significant increases in *COL1A1* (Fig 3F) and *FN* expression (Fig 3G) in aVICs compared to the untreated control. While the upregulation of *COL1A1* was observed at all three concentrations, the *FN* gene was only upregulated at the highest concentration - 600 µM. Notably, when both myofibroblastic (Fig 3B-C) and ECM (Fig 3F-G) genes were significantly upregulated, the gene expression levels surpassed those induced by treatment with TGF-β_1_, the most widely characterized profibrotic cytokine in the aortic valve [40,42] .

In summary, these findings suggest that TMAO promotes profibrotic behavior in both qVICs and aVICs, driving increased ECM production, a critical hallmark in the early progression of valve disease [43].

### qVIC activation by TMAO is not sex-dependent

CAVD exhibits significant sex-based differences in its pathophysiology [43–45] partially driven by differences in the response to disease stimuli between males and females at the cellular level [46–48]. Thus, we investigated whether sex influenced the qVIC response to TMAO. Male and female qVICs were treated with TMAO (25 – 150 µM) and TGF-β_1_ (1 ng/mL). Treatment with TMAO at concentrations ranging from 75 µM to 150 µM led to a significant upregulation of the *ACTA2* (S5A Fig) and *SM22* (S5B Fig) genes in both male and female qVICs. Corresponding protein-level increases in α-SMA (Fig. 4A-B) and transgelin (S5C Fig) were also observed with TMAO treatment at concentrations of 75 µM to 150 µM for both sexes.

**Figure 4.**
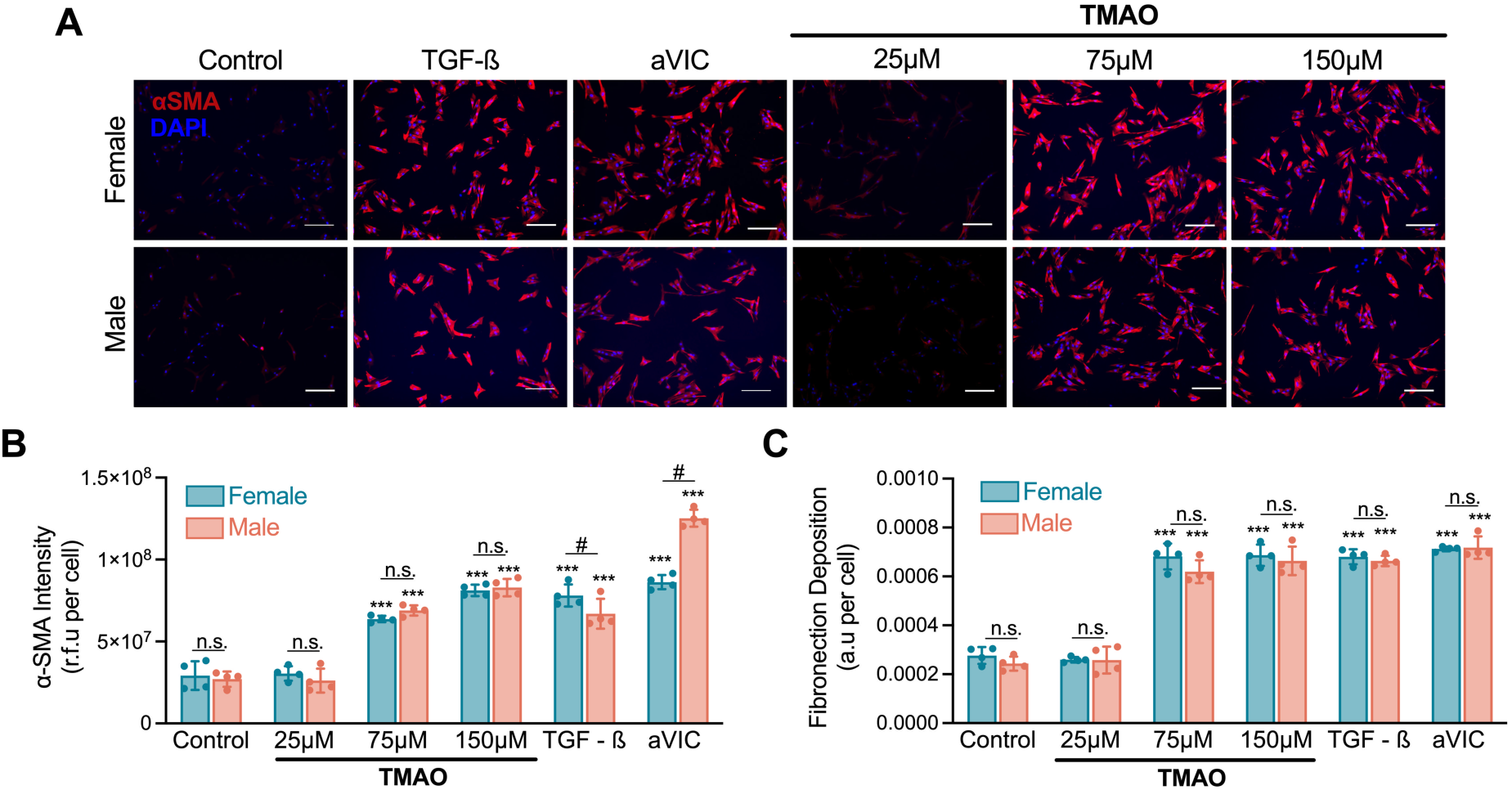
The activation of qVICs by TMAO does not depend on sex. Female and male qVICs were generated and treated with TMAO (25-150µM) for 3 days. (A) Representative images of immunocytochemistry staining for αSMA (red). Cell nuclei are stained blue. Scale bars represent 200 µm. (B) Quantification of αSMA staining intensity in (A). (C) Analysis of fibronectin deposition via *in situ* ELISA. n = 3-5 replicates per condition. Two-way ANOVA was performed, followed by Tukey’s multiple comparisons test. **p < 0.005, *p < 0.001 compared to the untreated control for the corresponding sex. n.s. denotes no statistically significant difference between male and female VICs for that condition. #*p*<0.01 for comparison shown.

For most treatment conditions, no sex-related differences were observed in the gene expression levels of these markers. However, for qVICs treated with 75 µM TMAO, there was a significantly higher upregulation of the *ACTA2* gene in the female cells compared to male VICs (S5A Fig), with the opposite effect for *SM22* (S5B Fig). No sex-based differences were detected in the expression of either α-SMA (Fig 4A-B) or transgelin (S5C Fig) at the protein level. Similar results were observed for proliferation (S5D Fig), were both male and female VICs proliferated at statistically higher rates after exposure to TMAO compared to the control, with no sex-based differences observed for any treatment condition.

As observed previously for female qVICs, male qVICs treated with TMAO concentrations equal or higher than 75 µM resulted in the upregulation of the ECM-related genes *FN* and *COL1A1* (S4E-F Fig). We also observed increased fibronectin deposition by qVICs of both sexes at these concentrations (Fig 4C). Consistent with our prior data, TMAO had no effect on collagen deposition (S4G Fig). No statistically significant differences in the response to TMAO were observed between male and female qVICs for any of these ECM-related end points.

These results demonstrate that TMAO treatment drives myofibroblastic activation and ECM production in both female and male qVICs, with minimal sex-based differences in the cellular response to this metabolite under the tested experimental conditions.

### TMAO triggers oxidative and ER stress in qVICs

Next, we explored the effects of TMAO on qVIC metabolism. Reactive oxygen species (ROS) have been implicated in both the progression of CAVD [49] and the cellular response to TMAO in other cardiovascular contexts [50]. Therefore, we assessed the production of ROS after treating female qVICs with TMAO for 3 days. TMAO led to an increase in intracellular ROS production at 75 µM - 150 µM concentrations compared to the untreated control (Fig 5A). These elevated ROS levels were also observed in the cell culture media (Fig 5B). Similar increases in ROS were observed in aVICs and after treatment with TGF-ß_1_.

**Figure 5.**
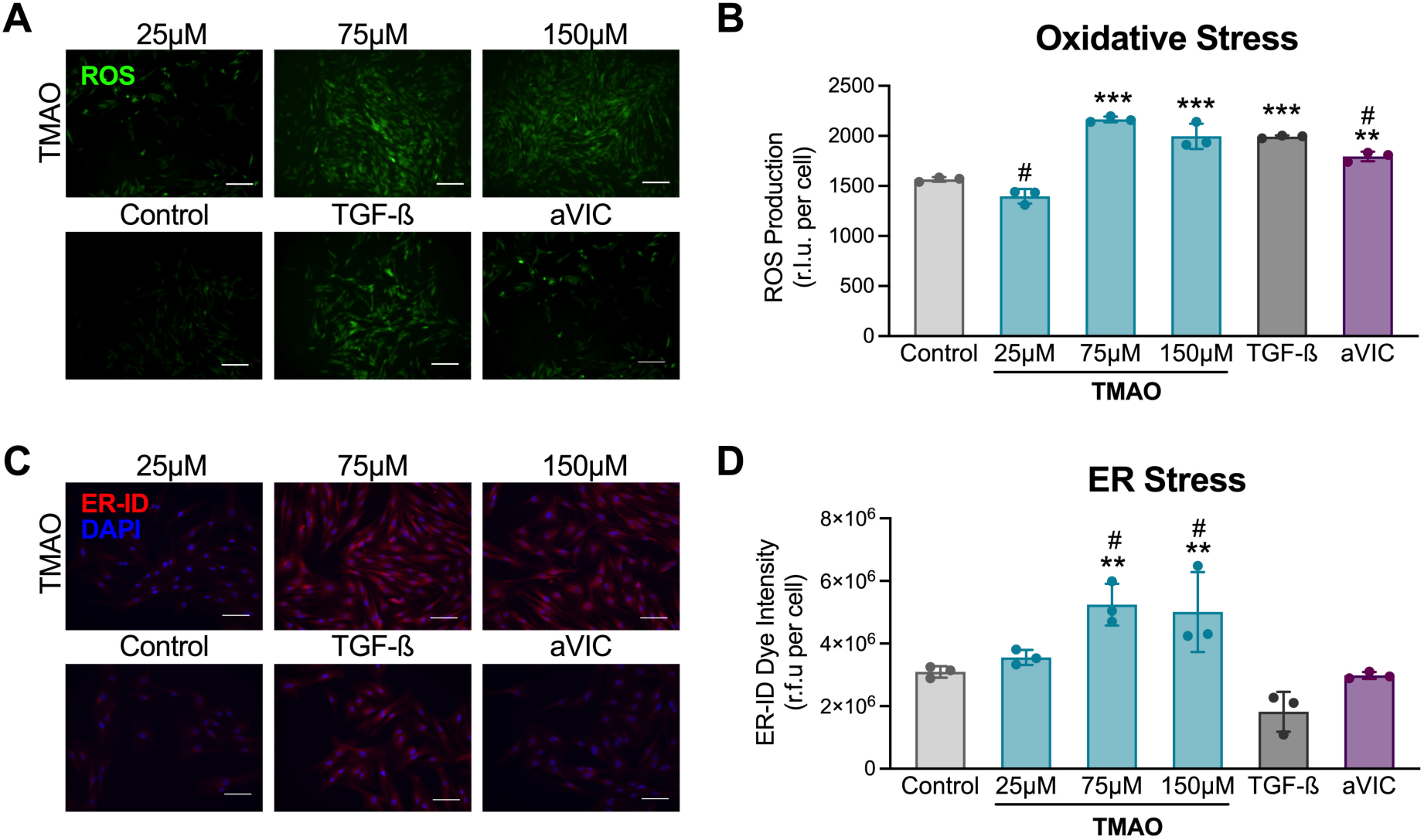
TMAO induces ROS production and ER stress. Female qVICs were treated with TMAO (25-150 µM) for 3 days. (A) Representative fluorescent images of intracellular 2’,7’-dichlorofluorescin diacetate (green), indicative of ROS. Scale bars represent 200 µm. (B) Quantification of the ROS-Glo^TM^ luminescent substrate in the culture media normalized to cell number. (C) Representative fluorescence images of the ER-ID Red dye, indicative of ER stress. Cell nuclei are stained blue. Scale bars represent 100 µm. (D) Quantification of fluorescence intensity in (C). n = 3-5 replicates per condition. One-way ANOVA followed by Tukey’s multiple comparisons test. \*\**p*<0.005, \*\*\**p*<0.001 compared with the control. #*p*<0.05 compared to treatment with TGF-ß_1_.

Because TMAO has also been reported to drive endoplasmic reticulum (ER) stress [51], we stained TMAO-treated qVICs with an ER-ID® Dye (Enzo Life Sciences). The detected intensity of this dye is proportional to the levels of ER stress in the stained cells [52]. Treatment with TMAO (75 µM) led to a statistically significant increase in ER stress compared to the untreated control (Fig 5C-D). No increase in ER stress was observed in qVICs treated with TGF-β_1_ or in the aVICs controls, suggesting this effect is unique to TMAO. These results establish that TMAO leads to metabolic dysfunction in qVICs through both oxidative and ER stress.

### TMAO drives qVIC activation through the PERK pathway

Finally, we sought to identify the specific molecular mechanisms through which TMAO activates qVICs. Chen *et al.* have previously demonstrated that TMAO drives ER stress and metabolic dysfunction in hepatocytes by directly binding and activating the endoplasmic reticulum stress kinase (PERK) [53]. Hence, we explored the role of the PERK pathway in the context of qVIC activation by TMAO. Female qVICs were pre-treated with a PERK inhibitor, GSK2656157, for 1 hour prior to treatment with either TMAO (150 µM) or TGF-β_1_ (1 ng/ml). GSK2656157 is a selective competitive inhibitor of PERK that blocks its activity by binding to the PERK kinase domain, thereby inhibiting its phosphorylation [54]. PERK inhibition successfully blocked the increase in ER stress previously observed after treatment with TMAO, while having no effect on the untreated or TGF-β controls (Fig 6A-B). Similarly, qVICs treated with the inhibitor did not exhibit an increase in ROS production after exposure to TMAO (Fig 6C). Inhibition of the PERK pathway had no effect on ROS production for qVICs treated with TGF-β_1_ (Fig 6C), suggesting that these two stimuli induce oxidative stress through distinct molecular pathways.

**Figure 6.**
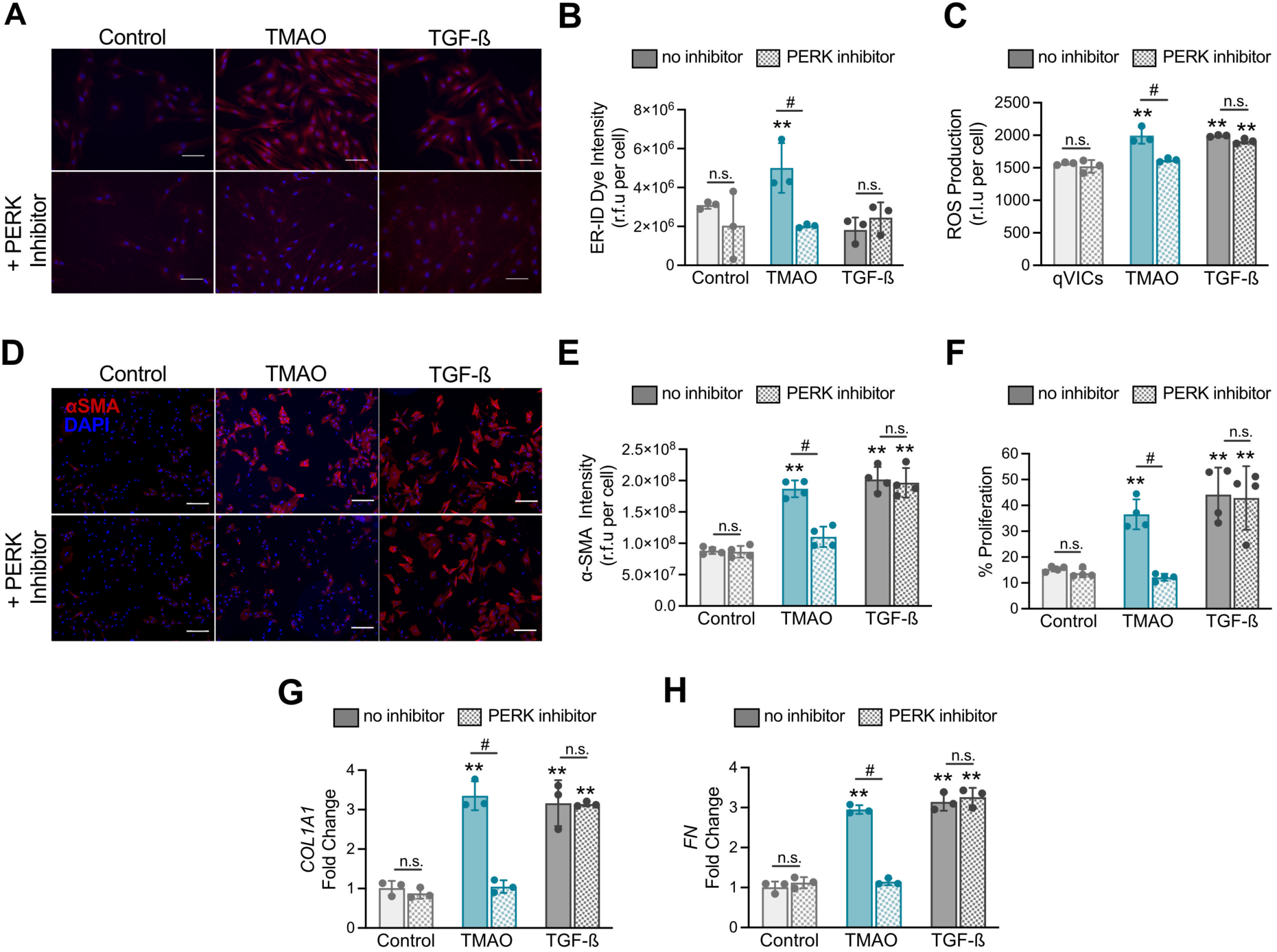
PERK inhibition mitigates the effects of TMAO. Female qVICs were pre-treated with GSK2656157 a PERK inhibitor, for 1 hour prior to treatment with TMAO. (A) Representative fluorescence images of the ER-ID Red dye, indicative of ER stress, at day 3. Cell nuclei are stained blue. Scale bars represent 100 µm. (B) Quantification of fluorescence intensity in (A). (C) Representative images of immunocytochemistry staining for αSMA (red) after 3 days of TMAO treatment. Cell nuclei are stained blue. Scale bars represent 200 µm. (D) Quantification of αSMA staining intensity in (B). (E) Quantification of the percentage of EdU-positive proliferating cells at day 3. (D) Quantification of ROS in the culture media through the ROS-Glo^TM^ luminescent substrate normalized to cell number. (G-H) Gene expression analysis of (G) COL1A1 and (H) *FN.* n = 3-5 replicates per condition. Two-way ANOVA, followed by Tukey’s multiple comparisons test. **p<0.005, ***p<0.001 compared with the control. n.s. denotes no statistically significant differences found between samples treated with and without the PERK inhibitor. #*p*<0.001 indicates a significant difference between samples treated with the PERK inhibitor and those without.

Having established that the GSK2656157 inhibitor successfully prevented the metabolic dysfunction induced by TMAO, we explored whether it also blocked the qVIC transition to an activated myofibroblastic phenotype. PERK inhibition significantly reduced the expression of both α-SMA (Fig 6D-E) and transgelin (S6A-B Fig) at the gene (S6C-D) and protein levels in qVICs treated with TMAO, effectively suppressing the transition to a myofibroblastic phenotype. Similar effects were observed for proliferation rates (Fig 6F). Treatment with the inhibitor had no significant effect on apoptosis (S6E Fig).

We also evaluated the role of PERK in regulating the pro-fibrotic response to TMAO. qVICs pre-treated with the PERK inhibitor did not upregulate the expression of *COL1A1* (Fig 6G) and *FN* (Fig 6H) upon treatment with TMAO. For all phenotypic outcomes, TGF-β_1_-mediated qVIC activation remained unaffected by PERK inhibition (Fig 6C-H). Moreover, blocking of the TGF-β_1_ receptor with the SB431542 inhibitor, did not interfere with the effects of TMAO on qVICs (SFig 7). Overall, these findings support the hypothesis that TMAO induces qVIC activation through ER stress and signaling via the PERK pathway.

## DISCUSSION

The gut microbiome is increasingly recognized as an important contributor to heart health [55–58]. More specifically, microbiome-derived metabolites like TMAO have been implicated in the initiation and progression of multiple cardiovascular diseases [59–63]. Patients with aortic valve stenosis exhibit significantly elevated serum TMAO levels compared to healthy controls, suggesting a relationship between this metabolite and valve disease [32]. However, the specific mechanisms by which TMAO contributes to CAVD are not fully understood. In the current study, we demonstrate that treatment with TMAO induces VIC activation towards a profibrotic myofibroblastic phenotype through molecular pathways specific to endoplasmic reticulum stress. Because VIC activation is a critical early step in the progression of disease [64–66], these results suggest that TMAO may play an important role in the initiation of CAVD.

Prior research demonstrated that TMAO stimulates osteogenic differentiation [31] and increased αSMA expression [30] in conventionally cultured human aVICs. Here, we expand upon this work by focusing on VICs of a quiescent phenotype and the critical early myofibroblastic transformation that precedes the osteogenic differentiation characteristic of later disease stages [43,64]. We found that treatment of porcine qVICs with TMAO at concentrations greater than 75μM led to an upregulation of myofibroblastic markers and increased proliferation, indicating the transition to an activated myofibroblast-like phenotype. Li *et al.* had previously observed that treatment of human aVICs with higher concentrations (200μM) of TMAO leads to the upregulation of osteoblastic markers like alkaline phosphatase and bone morphogenic protein 4 [31]. It is therefore possible that TMAO may exert differential effects on VICs depending on both phenotype and concentration.

In support of this hypothesis, quiescent VICs responded to TMAO treatment at concentrations 4 to 10-fold lower than activated VICs in our experiments. This result aligns with prior work demonstrating that qVICs are significantly more sensitive than aVICs to treatment with TGF-β_1_ [38]. Clinically, a mean difference of 2.2 µM TMAO separates patients with stroke from the control group, indicating that even modest differences in TMAO levels can have pathological significance [67]. The differences in the response to profibrotic stimuli between qVICs and aVICs underscore the importance of designing *in vitro* models that include cells with a physiological phenotype appropriate to the research question. In this study, the intentional use of VICs in a quiescent state provided valuable insights into the early cellular responses to TMAO.

In the kidney, heart, and liver, TMAO has been implicated in fibrosis by driving not only fibroblast activation but also ECM deposition [68–71]. This appears to also be the case in the aortic valve. Recently, Xiong *et al.* demonstrated that exposing conventionally cultured human VICs to TMAO increases collagen deposition *in vitro* and *in vivo* [30]. Our study builds upon these findings by showing that TMAO upregulates ECM production in both quiescent and activated VICs. More specifically, we observed the upregulation of fibronectin secretion at both the gene and protein level for both VIC phenotypes. On the other hand, collagen I was upregulated at the gene but not the protein level. This difference between *COL1A1* gene expression levels and collagen I deposition may be attributed to several factors, including post-translational modifications, increased matrix metalloproteinase activity, or impaired collagen secretion and assembly [72,73]. It may also be explained by a potential limitation of the *in-situ* ELISA assay, where collagen protein levels may have reached saturation, thus limiting the detection of subtle changes in collagen deposition [74]. Alterations in the content and organization of the ECM significantly impact the mechanical and biological properties of valve leaflets [75,76]. Therefore, increases in ECM production driven by TMAO may directly promote the progression of valve disease and contribute to impaired valve function through leaflet thickening, increased stiffness, and the exacerbation of pathological cell behavior [77–82].

A key finding of our study is that TMAO induces qVIC activation through molecular mechanisms independent of TGF-β signaling, the most widely studied pathway for VIC activation [42,83,84]. Treatment of qVICs with both TMAO and TGF-β_1_ significantly impacted VIC metabolism, with both stimuli resulting in a state of oxidative stress. However, only TMAO led to ER stress. Furthermore, inhibiting the PERK pathway did not affect TGF-β_1_-induced activation, nor did inhibition of the TGF-β receptor influence TMAO-driven activation. This indicates that TMAO and TGF-β_1_ activate qVICs through distinct molecular mechanisms despite driving similar outcomes—myofibroblastic transition, ECM production, and ROS generation.

We are not the first to link TMAO to increased ER stress. This metabolite has been proposed as a biomarker for pathogenic ER stress in the lung [85] and is known to induce ER stress in the kidney [69], liver [86], and heart [87]. ER stress usually activates all three pathways of the unfolded protein response: PERK, inositol-requiring enzyme 1 (IRE1), and activating transcription factor 6 (ATF6) [88]. In 2019, Chen *et al.* identified PERK as a receptor for TMAO, showing that TMAO directly binds to and selectively activates the PERK branch in renal cells [53]. In our study, blocking the PERK receptor effectively inhibited TMAO-induced qVIC myofibroblastic differentiation and ECM production, with previous studies observing similar effects in human VICs [30,31]. Collectively, this evidence indicates that TMAO also exerts its effects in the aortic valve through the PERK pathway. Given the past failures of lipid-lowering approaches [89,90], furthering the understanding of the PERK pathway and other metabolic pathways triggered by TMAO may lead to the identification of novel potential therapeutic or diagnostic markers.

Sex is a key factor and biological variable in CAVD, with men exhibiting a higher prevalence of calcification [91–94], and women displaying greater fibrotic remodeling [92]. Cellular-scale sex differences in VIC behavior have also been observed *in vitro* [47,95,96]. These sex-specific manifestations suggest that the mechanisms driving CAVD could differ between males and females. Here, we found that TMAO activates both male and female qVICs with no statistically significant differences in the response between sexes, suggesting that sex may not be a major biological variable influencing the response to this metabolite. However, it is important to note that these cells were cultured on a stiff 2D substrate. Prior research has shown that male and female VICs exhibit distinct behaviors when cultured in 3D substrates [48,97,98]. For example, myofibroblastic activation is higher in female than male VICs on soft hydrogels that resemble a healthy valve microenvironment, with an even greater sex difference in activation observed in stiff substrates [48]. Thus, future studies leveraging these 3D culture systems will be necessary to definitively establish whether the VIC response to TMAO varies depending on cellular sex.

There are additional important considerations for the interpretation of our results. First, we used porcine VICs for our experiments. This a common practice in the field due to their similarities with human cells [99–101], the absence of a human cell line, and the challenges associated with procuring human tissue [99]. However, species-specific differences may affect cellular responses. For example, human VICs are less prone to spontaneous *in vitro* activation than porcine VICs [38]. Nonetheless, others have demonstrated that TMAO also affects the phenotype of human VICS *in vitro* [28,30], suggesting that our findings are applicable across species. Second, despite the increased sensitivity of qVICs compared to conventionally cultured cells, our study utilized TMAO at concentrations higher than those present physiologically. Serum TMAO levels hover around 1-2μM and 2-7μM for healthy and aortic valve stenosis patients, respectively [32]. Considering the chronic nature of valve disease and most cardiovascular conditions, it is possible that prolonged exposure to this metabolite is necessary to observe effects at low TMAO concentrations. Clark-Greuel *et al.* demonstrated that prolonged TGF-β_1_ exposure significantly influenced VIC behavior, leading to increased calcium deposition and calcification observed over 14 days compared to shorter exposures of 3 or 7 days [102]. This highlights the potential for time-dependent changes in VIC responses to profibrotic stimuli. Future experiments focusing on the impact of long-term exposure to lower, clinically relevant TMAO concentrations may lead to the emergence of additional pathological phenotypes or calcification markers that would not manifest in shorter studies.

In summary, our study provides critical insights into the role of TMAO in the early stages of CAVD, demonstrating its ability to activate qVICs towards a profibrotic myofibroblastic phenotype via the PERK pathway, a key regulator of ER stress and oxidative imbalance. By leveraging our culture strategy to maintain VICs in their quiescent phenotype, we were able to identify that quiescent VICs respond to TMAO treatment at much lower concentrations than conventionally cultured activated VICs. This observation emphasizes the importance of developing physiologically relevant *in vitro* models that mimic the conditions seen in healthy and early disease valves. Overall, our results contribute to the growing body of knowledge connecting dietary patterns and the gut microbiome to the regulation of cardiovascular health. Understanding the contributions of TMAO and other gut metabolites to the early stages of valve disease may pave the way for the development of preventative strategies to delay the progression of CAVD or targeted treatment alternatives beyond valve replacement. Moreover, these insights also emphasize the potential for dietary interventions to mitigate cardiovascular and metabolic disease risk.

## ACKNOWLEDGEMENTS

We thank Dr. Abisambra from the Department of Neuroscience at the University of Florida for valuable discussions related to the PERK signaling pathway. We also thank Karen Mancera Azamar and Zahra Mohammadalizadeh for their assistance in experiment planning. Icons for the figures were created using BioRender©.

## AUTHOR CONTRIBUTIONS

Conceptualization and methodology: S.S. and A.M.P; investigation: S.S., S.D.S. and T.K.; formal analysis: S.S., S.D.S., and T.K., visualization and writing – original draft: S.S.; writing – review & editing: S.S. and A.M.P.; supervision and funding acquisition: A.M.P.

## REFERENCES

1. Yu J, Wang Z, Bao Q, Lei S, You Y, Yin Z, et al. Global burden of calcific aortic valve disease and attributable risk factors from 1990 to 2019. Front Cardiovasc Med. 2022;9: 1003233. doi:10.3389/fcvm.2022.1003233

2. Santangelo G, Bursi F, Faggiano A, Moscardelli S, Simeoli P, Guazzi M, et al. The Global Burden of Valvular Heart Disease: From Clinical Epidemiology to Management. JCM. 2023;12: 2178. doi:10.3390/jcm12062178

3. Kumar V, Sandhu GS, Harper CM, Ting HH, Rihal CS. Transcatheter Aortic Valve Replacement Programs: Clinical Outcomes and Developments. Journal of the American Heart Association. 2020;9: e015921. doi:10.1161/JAHA.120.015921

4. Witkowski M, Weeks TL, Hazen SL. Gut Microbiota and Cardiovascular Disease. Circulation Research. 2020;127: 553–570. doi:10.1161/CIRCRESAHA.120.316242

5. Trøseid M, Andersen GØ, Broch K, Hov JR. The gut microbiome in coronary artery disease and heart failure: Current knowledge and future directions. EBioMedicine. 2020;52: 102649. doi:10.1016/j.ebiom.2020.102649

6. Jie Z, Xia H, Zhong S-L, Feng Q, Li S, Liang S, et al. The gut microbiome in atherosclerotic cardiovascular disease. Nat Commun. 2017;8: 845. doi:10.1038/s41467-017-00900-1

7. Chen X, Zhang H, Ren S, Ding Y, Remex NS, Bhuiyan MdS, et al. Gut microbiota and microbiota-derived metabolites in cardiovascular diseases. Chinese Medical Journal. 2023;136: 2269–2284. doi:10.1097/CM9.0000000000002206

8. Hemmati M, Kashanipoor S, Mazaheri P, Alibabaei F, Babaeizad A, Asli S, et al. Importance of gut microbiota metabolites in the development of cardiovascular diseases (CVD). Life Sciences. 2023;329: 121947. doi:10.1016/j.lfs.2023.121947

9. Zhen J, Zhou Z, He M, Han H-X, Lv E-H, Wen P-B, et al. The gut microbial metabolite trimethylamine N-oxide and cardiovascular diseases. Front Endocrinol. 2023;14: 1085041. doi:10.3389/fendo.2023.1085041

10. Koeth RA, Wang Z, Levison BS, Buffa JA, Org E, Sheehy BT, et al. Intestinal microbiota metabolism of l-carnitine, a nutrient in red meat, promotes atherosclerosis. Nat Med. 2013;19: 576–585. doi:10.1038/nm.3145

11. Jing L, Zhang H, Xiang Q, Hu H, Zhai C, Xu S, et al. Role of Trimethylamine N-Oxide in Heart Failure. RCM. 2024;25: 240. doi:10.31083/j.rcm2507240

12. Wang Z, Klipfell E, Bennett BJ, Koeth R, Levison BS, DuGar B, et al. Gut flora metabolism of phosphatidylcholine promotes cardiovascular disease. Nature. 2011;472: 57–63. doi:10.1038/nature09922

13. Rath S, Heidrich B, Pieper DH, Vital M. Uncovering the trimethylamine-producing bacteria of the human gut microbiota. Microbiome. 2017;5: 54. doi:10.1186/s40168-017-0271-9

14. Carnitine metabolism to trimethylamine by an unusual Rieske-type oxygenase from human microbiota. [cited 21 Jan 2025]. doi:10.1073/pnas.1316569111

15. Zhu W, Buffa JA, Wang Z, Warrier M, Schugar R, Shih DM, et al. Flavin monooxygenase 3, the host hepatic enzyme in the metaorganismal trimethylamine N-oxide-generating pathway, modulates platelet responsiveness and thrombosis risk. J Thromb Haemost. 2018;16: 1857–1872. doi:10.1111/jth.14234

16. Bennett BJ, de Aguiar Vallim TQ, Wang Z, Shih DM, Meng Y, Gregory J, et al. Trimethylamine-N-Oxide, a Metabolite Associated with Atherosclerosis, Exhibits Complex Genetic and Dietary Regulation. Cell Metab. 2013;17: 49–60. doi:10.1016/j.cmet.2012.12.011

17. Treacy E. Mutations of the flavin-containing monooxygenase gene (FMO3) cause trimethylaminuria, a defect in detoxication. Human Molecular Genetics. 1998;7: 839–845. doi:10.1093/hmg/7.5.839

18. Heianza Y, Ma W, Manson JE, Rexrode KM, Qi L. Gut Microbiota Metabolites and Risk of Major Adverse Cardiovascular Disease Events and Death: A Systematic Review and Meta-Analysis of Prospective Studies. Journal of the American Heart Association. 2017;6: e004947. doi:10.1161/JAHA.116.004947

19. Heianza Y, Ma W, DiDonato JA, Sun Q, Rimm EB, Hu FB, et al. Long-Term Changes in Gut Microbial Metabolite Trimethylamine N-Oxide and Coronary Heart Disease Risk. Journal of the American College of Cardiology. 2020;75: 763–772. doi:10.1016/j.jacc.2019.11.060

20. Naghipour S, Cox AJ, Fisher JJ, Plan M, Stark T, West N, et al. Circulating TMAO, the gut microbiome and cardiometabolic disease risk: an exploration in key precursor disorders. Diabetol Metab Syndr. 2024;16: 133. doi:10.1186/s13098-024-01368-y

21. Ge X, Zheng L, Zhuang R, Yu P, Xu Z, Liu G, et al. The Gut Microbial Metabolite Trimethylamine N-Oxide and Hypertension Risk: A Systematic Review and Dose– Response Meta-analysis. Advances in Nutrition. 2020;11: 66–76. doi:10.1093/advances/nmz064

22. Lee Y, Nemet I, Wang Z, Lai HTM, De Oliveira Otto MC, Lemaitre RN, et al. Longitudinal Plasma Measures of Trimethylamine N-Oxide and Risk of Atherosclerotic Cardiovascular Disease Events in Community-Based Older Adults. JAHA. 2021;10: e020646. doi:10.1161/JAHA.120.020646

23. Wu P, Chen J, Chen J, Tao J, Wu S, Xu G, et al. Trimethylamine N-oxide promotes apoE−/− mice atherosclerosis by inducing vascular endothelial cell pyroptosis via the SDHB/ROS pathway. Journal of Cellular Physiology. 2020;235: 6582–6591. doi:10.1002/jcp.29518

24. Cheng X, Qiu X, Liu Y, Yuan C, Yang X. Trimethylamine N-oxide promotes tissue factor expression and activity in vascular endothelial cells: A new link between trimethylamine N-oxide and atherosclerotic thrombosis. Thrombosis Research. 2019;177: 110–116. doi:10.1016/j.thromres.2019.02.028

25. Catar R, Chen L, Zhao H, Wu D, Kamhieh-Milz J, Lücht C, et al. Native and Oxidized Low-Density Lipoproteins Increase the Expression of the LDL Receptor and the LOX-1 Receptor, Respectively, in Arterial Endothelial Cells. Cells. 2022;11: 204. doi:10.3390/cells11020204

26. Koh Y-C, Li S, Chen P-Y, Wu J-C, Kalyanam N, Ho C-T, et al. Prevention of Vascular Inflammation by Pterostilbene via Trimethylamine-N-Oxide Reduction and Mechanism of Microbiota Regulation. Molecular Nutrition & Food Research. 2019;63: 1900514. doi:10.1002/mnfr.201900514

27. Geng J, Yang C, Wang B, Zhang X, Hu T, Gu Y, et al. Trimethylamine N-oxide promotes atherosclerosis via CD36-dependent MAPK/JNK pathway. Biomedicine & Pharmacotherapy. 2018;97: 941–947. doi:10.1016/j.biopha.2017.11.016

28. Li X, Geng J, Zhao J, Ni Q, Zhao C, Zheng Y, et al. Trimethylamine N-Oxide Exacerbates Cardiac Fibrosis via Activating the NLRP3 Inflammasome. Front Physiol. 2019;10. doi:10.3389/fphys.2019.00866

29. Nayak G, Dimitriadis K, Pyrpyris N, Manti M, Kamperidis N, Kamperidis V, et al. Gut Microbiome and Its Role in Valvular Heart Disease: Not a “Gutted” Relationship. Life. 2024;14: 527. doi:10.3390/life14040527

30. Xiong Z, Li J, Huang R, Zhou H, Xu X, Zhang S, et al. The gut microbe-derived metabolite trimethylamine-N-oxide induces aortic valve fibrosis via PERK/ATF-4 and IRE-1α/XBP-1s signaling in vitro and in vivo. Atherosclerosis. 2024;391: 117431. doi:10.1016/j.atherosclerosis.2023.117431

31. Li J, Zeng Q, Xiong Z, Xian G, Liu Z, Zhan Q, et al. Trimethylamine N-oxide induces osteogenic responses in human aortic valve interstitial cells *in vitro* and aggravates aortic valve lesions in mice. Cardiovascular Research. 2022;118: 2018– 2030. doi:10.1093/cvr/cvab243

32. Guo Y, Xu S, Zhan H, Chen H, Hu P, Zhou D, et al. Trimethylamine N-Oxide Levels Are Associated with Severe Aortic Stenosis and Predict Long-Term Adverse Outcome. JCM. 2023;12: 407. doi:10.3390/jcm12020407

33. Rutkovskiy A, Malashicheva A, Sullivan G, Bogdanova M, Kostareva A, Stensløkken K, et al. Valve Interstitial Cells: The Key to Understanding the Pathophysiology of Heart Valve Calcification. JAHA. 2017;6: e006339. doi:10.1161/JAHA.117.006339

34. Chester AH, Taylor PM. Molecular and functional characteristics of heart-valve interstitial cells. Phil Trans R Soc B. 2007;362: 1437–1443. doi:10.1098/rstb.2007.2126

35. Liu AC, Joag VR, Gotlieb AI. The Emerging Role of Valve Interstitial Cell Phenotypes in Regulating Heart Valve Pathobiology. The American Journal of Pathology. 2007;171: 1407–1418. doi:10.2353/ajpath.2007.070251

36. Ali MS, Deb N, Wang X, Rahman M, Christopher GF, Lacerda CMR. Correlation between valvular interstitial cell morphology and phenotypes: A novel way to detect activation. Tissue and Cell. 2018;54: 38–46. doi:10.1016/j.tice.2018.07.004

37. Ma H, Killaars AR, DelRio FW, Yang C, Anseth KS. Myofibroblastic activation of valvular interstitial cells is modulated by spatial variations in matrix elasticity and its organization. Biomaterials. 2017;131: 131–144. doi:10.1016/j.biomaterials.2017.03.040

38. Porras AM, van Engeland NCA, Marchbanks E, McCormack A, Bouten CVC, Yacoub MH, et al. Robust Generation of Quiescent Porcine Valvular Interstitial Cell Cultures. Journal of the American Heart Association. 2017;6: e005041. doi:10.1161/JAHA.116.005041

39. Frangogiannis NG. Transforming growth factor–β in tissue fibrosis. Journal of Experimental Medicine. 2020;217: e20190103. doi:10.1084/jem.20190103

40. Hirai H, Yang B, Garcia-Barrio MT, Rom O, Ma PX, Zhang J, et al. Direct Reprogramming of Fibroblasts Into Smooth Muscle-Like Cells With Defined Transcription Factors—Brief Report. ATVB. 2018;38: 2191–2197. doi:10.1161/ATVBAHA.118.310870

41. Chmelova M, Androvic P, Kirdajova D, Tureckova J, Kriska J, Valihrach L, et al. A view of the genetic and proteomic profile of extracellular matrix molecules in aging and stroke. Front Cell Neurosci. 2023;17. doi:10.3389/fncel.2023.1296455

42. Walker GA, Masters KS, Shah DN, Anseth KS, Leinwand LA. Valvular Myofibroblast Activation by Transforming Growth Factor-β. Circulation Research. 2004;95: 253–260. doi:10.1161/01.RES.0000136520.07995.aa

43. Rajamannan NM, Evans FJ, Aikawa E, Grande-Allen KJ, Demer LL, Heistad DD, et al. Calcific Aortic Valve Disease: Not Simply a Degenerative Process. Circulation. 2011;124: 1783–1791. doi:10.1161/CIRCULATIONAHA.110.006767

44. Le Nezet E, Marqueze-Pouey C, Guisle I, Clavel M-A. Molecular Features of Calcific Aortic Stenosis in Female and Male Patients. CJC Open. 2024;6: 1125– 1137. doi:10.1016/j.cjco.2024.06.002

45. Myasoedova VA, Massaiu I, Moschetta D, Chiesa M, Songia P, Valerio V, et al. Sex-Specific Cell Types and Molecular Pathways Indicate Fibro-Calcific Aortic Valve Stenosis. Front Immunol. 2022;13: 747714. doi:10.3389/fimmu.2022.747714

46. McCoy CM, Nicholas DQ, Masters KS. Sex-Related Differences in Gene Expression by Porcine Aortic Valvular Interstitial Cells. Aikawa E, editor. PLoS ONE. 2012;7: e39980. doi:10.1371/journal.pone.0039980

47. Simon LR, Scott AJ, Figueroa Rios L, Zembles J, Masters KS. Cellular-scale sex differences in extracellular matrix remodeling by valvular interstitial cells. Heart Vessels. 2023;38: 122–130. doi:10.1007/s00380-022-02164-2

48. Aguado BA, Walker CJ, Grim JC, Schroeder ME, Batan D, Vogt BJ, et al. Genes That Escape X Chromosome Inactivation Modulate Sex Differences in Valve Myofibroblasts. Circulation. 2022;145: 513–530. doi:10.1161/CIRCULATIONAHA.121.054108

49. Greenberg HZE, Zhao G, Shah AM, Zhang M. Role of oxidative stress in calcific aortic valve disease and its therapeutic implications. Cardiovascular Research. 2022;118: 1433–1451. doi:10.1093/cvr/cvab142

50. Saaoud F, Liu L, Xu K, Cueto R, Shao Y, Lu Y, et al. Aorta- and liver-generated TMAO enhances trained immunity for increased inflammation via ER stress/mitochondrial ROS/glycolysis pathways. JCI Insight. 2023;8. doi:10.1172/jci.insight.158183

51. Govindarajulu M, Pinky PD, Steinke I, Bloemer J, Ramesh S, Kariharan T, et al. Gut Metabolite TMAO Induces Synaptic Plasticity Deficits by Promoting Endoplasmic Reticulum Stress. Front Mol Neurosci. 2020;13. doi:10.3389/fnmol.2020.00138

52. Wang Y-Y, Lee K-T, Lim MC, Choi J-H. TRPV1 Antagonist DWP05195 Induces ER Stress-Dependent Apoptosis through the ROS-p38-CHOP Pathway in Human Ovarian Cancer Cells. Cancers (Basel). 2020;12: 1702. doi:10.3390/cancers12061702

53. Chen S, Henderson A, Petriello MC, Romano KA, Gearing M, Miao J, et al. Trimethylamine N-Oxide Binds and Activates PERK to Promote Metabolic Dysfunction. Cell Metabolism. 2019;30: 1141–1151.e5. doi:10.1016/j.cmet.2019.08.021

54. Axten JM, Romeril SP, Shu A, Ralph J, Medina JR, Feng Y, et al. Discovery of GSK2656157: An Optimized PERK Inhibitor Selected for Preclinical Development. ACS Med Chem Lett. 2013;4: 964–968. doi:10.1021/ml400228e

55. Masenga SK, Hamooya B, Hangoma J, Hayumbu V, Ertuglu LA, Ishimwe J, et al. Recent advances in modulation of cardiovascular diseases by the gut microbiota. J Hum Hypertens. 2022;36: 952–959. doi:10.1038/s41371-022-00698-6

56. Wang L, Wang S, Zhang Q, He C, Fu C, Wei Q. The role of the gut microbiota in health and cardiovascular diseases. Mol Biomed. 2022;3: 30. doi:10.1186/s43556-022-00091-2

57. Ahmad AF, Dwivedi G, O’Gara F, Caparros-Martin J, Ward NC. The gut microbiome and cardiovascular disease: current knowledge and clinical potential. American Journal of Physiology-Heart and Circulatory Physiology. 2019;317: H923– H938. doi:10.1152/ajpheart.00376.2019

58. Tang WHW, Kitai T, Hazen SL. Gut Microbiota in Cardiovascular Health and Disease. Circulation Research. 2017;120: 1183–1196. doi:10.1161/CIRCRESAHA.117.309715

59. Roncal C, Martínez-Aguilar E, Orbe J, Ravassa S, Fernandez-Montero A, Saenz-Pipaon G, et al. Trimethylamine-N-Oxide (TMAO) Predicts Cardiovascular Mortality in Peripheral Artery Disease. Sci Rep. 2019;9: 15580. doi:10.1038/s41598-019-52082-z

60. Trøseid M, Andersen GØ, Broch K, Hov JR. The gut microbiome in coronary artery disease and heart failure: Current knowledge and future directions. EBioMedicine. 2020;52: 102649. doi:10.1016/j.ebiom.2020.102649

61. Meng G, Zhou X, Wang M, Zhou L, Wang Z, Wang M, et al. Gut microbe-derived metabolite trimethylamine N-oxide activates the cardiac autonomic nervous system and facilitates ischemia-induced ventricular arrhythmia via two different pathways. eBioMedicine. 2019;44: 656–664. doi:10.1016/j.ebiom.2019.03.066

62. Nayak G, Dimitriadis K, Pyrpyris N, Manti M, Kamperidis N, Kamperidis V, et al. Gut Microbiome and Its Role in Valvular Heart Disease: Not a “Gutted” Relationship. Life. 2024;14: 527. doi:10.3390/life14040527

63. Karlin ET, Rush JE, Freeman LM. A pilot study investigating circulating trimethylamine N-oxide and its precursors in dogs with degenerative mitral valve disease with or without congestive heart failure. J Vet Intern Med. 2019;33: 46–53. doi:10.1111/jvim.15347

64. Monzack EL, Masters KS. Can valvular interstitial cells become true osteoblasts?: A side-by-side comparison. J Heart Valve Dis. 2011;20: 449–463.

65. Wu S, Kumar V, Xiao P, Kuss M, Lim JY, Guda C, et al. Age related extracellular matrix and interstitial cell phenotype in pulmonary valves. Sci Rep. 2020;10: 21338. doi:10.1038/s41598-020-78507-8

66. Ohukainen P, Ruskoaho H, Rysä J. Cellular Mechanisms of Valvular Thickening in Early and Intermediate Calcific Aortic Valve Disease. Curr Cardiol Rev. 2018;14: 264–271. doi:10.2174/1573403X14666180820151325

67. Farhangi MA, Vajdi M, Asghari-Jafarabadi M. Gut microbiota-associated metabolite trimethylamine N-Oxide and the risk of stroke: a systematic review and dose–response meta-analysis. Nutr J. 2020;19: 76. doi:10.1186/s12937-020-00592-2

68. Stefania K, Ashok KK, Geena PV, Katarina P, Isak D. TMAO enhances TNF-α mediated fibrosis and release of inflammatory mediators from renal fibroblasts. Sci Rep. 2024;14: 9070. doi:10.1038/s41598-024-58084-w

69. Kapetanaki S, Kumawat AK, Persson K, Demirel I. The Fibrotic Effects of TMAO on Human Renal Fibroblasts Is Mediated by NLRP3, Caspase-1 and the PERK/Akt/mTOR Pathway. IJMS. 2021;22: 11864. doi:10.3390/ijms222111864

70. Zhou D, Zhang J, Xiao C, Mo C, Ding B-S. Trimethylamine-N-oxide (TMAO) mediates the crosstalk between the gut microbiota and hepatic vascular niche to alleviate liver fibrosis in nonalcoholic steatohepatitis. Front Immunol. 2022;13: 964477. doi:10.3389/fimmu.2022.964477

71. Kim S-J, Bale S, Verma P, Wan Q, Ma F, Gudjonsson JE, et al. Gut microbe-derived metabolite trimethylamine N-oxide activates PERK to drive fibrogenic mesenchymal differentiation. iScience. 2022;25: 104669. doi:10.1016/j.isci.2022.104669

72. Myllyharju J. Intracellular Post-Translational Modifications of Collagens. In: Brinckmann J, Notbohm H, Müller PK, editors. Collagen: Primer in Structure, Processing and Assembly. Berlin, Heidelberg: Springer; 2005. pp. 115–147. doi:10.1007/b103821

73. Naomi R, Ridzuan PM, Bahari H. Current Insights into Collagen Type I. Polymers. 2021;13: 2642. doi:10.3390/polym13162642

74. Byun J-Y, Lee K-H, Park YJ, Song D-Y, Min Y-H, Kim D-M. Revisiting ELISA with *in situ* amplification of biomarkers to boost its sensitivity. Sensors and Actuators B: Chemical. 2025;423: 136780. doi:10.1016/j.snb.2024.136780

75. Chen J-H, Simmons CA. Cell–Matrix Interactions in the Pathobiology of Calcific Aortic Valve Disease: Critical Roles for Matricellular, Matricrine, and Matrix Mechanics Cues. Towler DA, editor. Circulation Research. 2011;108: 1510–1524. doi:10.1161/CIRCRESAHA.110.234237

76. Scott AJ, Simon LR, Hutson HN, Porras AM, Masters KS. Engineering the aortic valve extracellular matrix through stages of development, aging, and disease. Journal of Molecular and Cellular Cardiology. 2021;161: 1–8. doi:10.1016/j.yjmcc.2021.07.009

77. Duan B, Yin Z, Hockaday Kang L, Magin RL, Butcher JT. Active tissue stiffness modulation controls valve interstitial cell phenotype and osteogenic potential in 3D culture. Acta Biomaterialia. 2016;36: 42–54. doi:10.1016/j.actbio.2016.03.007

78. Hjortnaes J, Goettsch C, Hutcheson JD, Camci-Unal G, Lax L, Scherer K, et al. Simulation of early calcific aortic valve disease in a 3D platform: A role for myofibroblast differentiation. Journal of Molecular and Cellular Cardiology. 2016;94: 13–20. doi:10.1016/j.yjmcc.2016.03.004

79. Rodriguez KJ, Piechura LM, Masters KS. Regulation of valvular interstitial cell phenotype and function by hyaluronic acid in 2-D and 3-D culture environments. Matrix Biology. 2011;30: 70–82. doi:10.1016/j.matbio.2010.09.001

80. Rodriguez KJ, Masters KS. Regulation of valvular interstitial cell calcification by components of the extracellular matrix. Journal of Biomedical Materials Research Part A. 2009;90A: 1043–1053. doi:10.1002/jbm.a.32187

81. Porras AM, Westlund JA, Evans AD, Masters KS. Creation of disease-inspired biomaterial environments to mimic pathological events in early calcific aortic valve disease. Proceedings of the National Academy of Sciences. 2018; 201704637. doi:10.1073/pnas.1704637115

82. Monroe MN, Nikonowicz RC, Grande-Allen KJ. Heterogeneous multi-laminar tissue constructs as a platform to evaluate aortic valve matrix-dependent pathogenicity. Acta Biomaterialia. 2019;97: 420–427. doi:10.1016/j.actbio.2019.07.046

83. Jian B, Narula N, Li Q, Iii ERM, Levy RJ. Progression of Aortic Valve Stenosis: TGF-␤1 is Present in Calcified Aortic Valve Cusps and Promotes Aortic Valve Interstitial Cell Calcification Via Apoptosis. Ann Thorac Surg.

84. Cushing MC, Liao J-T, Anseth KS. Activation of valvular interstitial cells is mediated by transforming growth factor-β1 interactions with matrix molecules. Matrix Biology. 2005;24: 428–437. doi:10.1016/j.matbio.2005.06.007

85. Wang Z, Ma P, Wang Y, Hou B, Zhou C, Tian H, et al. Untargeted metabolomics and transcriptomics identified glutathione metabolism disturbance and PCS and TMAO as potential biomarkers for ER stress in lung. Sci Rep. 2021;11: 14680. doi:10.1038/s41598-021-92779-8

86. Yang B, Tang G, Wang M, Ni Y, Tong J, Hu C, et al. Trimethylamine N-oxide induces non-alcoholic fatty liver disease by activating the PERK. Toxicology Letters. 2024;400: 93–103. doi:10.1016/j.toxlet.2024.08.009

87. Cheng T-Y, Lee T-W, Li S-J, Lee T-I, Chen Y-C, Kao Y-H, et al. Short-chain fatty acid butyrate against TMAO activating endoplasmic-reticulum stress and PERK/IRE1-axis with reducing atrial arrhythmia. Journal of Advanced Research. 2024 [cited 29 Jan 2025]. doi:10.1016/j.jare.2024.08.009

88. Chen X, Shi C, He M, Xiong S, Xia X. Endoplasmic reticulum stress: molecular mechanism and therapeutic targets. Sig Transduct Target Ther. 2023;8: 352. doi:10.1038/s41392-023-01570-w

89. Myasoedova VA, Ravani AL, Frigerio B, Valerio V, Moschetta D, Songia P, et al. Novel pharmacological targets for calcific aortic valve disease: Prevention and treatments. Pharmacological Research. 2018;136: 74–82. doi:10.1016/j.phrs.2018.08.020

90. Moncla L-HM, Briend M, Bossé Y, Mathieu P. Calcific aortic valve disease: mechanisms, prevention and treatment. Nat Rev Cardiol. 2023;20: 546–559. doi:10.1038/s41569-023-00845-7

91. Ward LJ, Laucyte-Cibulskiene A, Hernandez L, Ripsweden J, Pilote L, Norris CM, et al. Coronary artery calcification and aortic valve calcification in patients with kidney failure: a sex-disaggregated study. Biology of Sex Differences. 2023;14: 48. doi:10.1186/s13293-023-00530-x

92. Voisine M, Hervault M, Shen M, Boilard A, Filion B, Rosa M, et al. Age, Sex, and Valve Phenotype Differences in Fibro-Calcific Remodeling of Calcified Aortic Valve. JAHA. 2020;9: e015610. doi:10.1161/JAHA.119.015610

93. Simard L, Côté N, Dagenais F, Mathieu P, Couture C, Trahan S, et al. Sex-Related Discordance Between Aortic Valve Calcification and Hemodynamic Severity of Aortic Stenosis. Circulation Research. 2017;120: 681–691. doi:10.1161/CIRCRESAHA.116.309306

94. Aggarwal SR, Clavel M-A, Messika-Zeitoun D, Cueff C, Malouf J, Araoz PA, et al. Sex Differences in Aortic Valve Calcification Measured by Multidetector Computed Tomography in Aortic Stenosis. Circ: Cardiovascular Imaging. 2013;6: 40–47. doi:10.1161/CIRCIMAGING.112.980052

95. Masjedi S, Lei Y, Patel J, Ferdous Z. Sex-related differences in matrix remodeling and early osteogenic markers in aortic valvular interstitial cells. Heart Vessels. 2017;32: 217–228. doi:10.1007/s00380-016-0909-8

96. Nelson V, Patil V, Simon LR, Schmidt K, McCoy CM, Masters KS. Angiogenic Secretion Profile of Valvular Interstitial Cells Varies With Cellular Sex and Phenotype. Front Cardiovasc Med. 2021;8: 736303. doi:10.3389/fcvm.2021.736303

97. Félix Vélez NE, Tu K, Guo P, Reeves RR, Aguado BA. Secreted cytokines from inflammatory macrophages modulate sex differences in valvular interstitial cells on hydrogel biomaterials. 2024. doi:10.1101/2024.11.15.623805

98. Vogt BJ, Peters DK, Anseth KS, Aguado BA. Inflammatory serum factors from aortic valve stenosis patients modulate sex differences in valvular myofibroblast activation and osteoblast-like differentiation. Biomater Sci. 2022;10: 6341–6353. doi:10.1039/D2BM00844K

99. Nehl D, Goody PR, Maus K, Pfeifer A, Aikawa E, Bakthiary F, et al. Human and porcine aortic valve endothelial and interstitial cell isolation and characterization. Front Cardiovasc Med. 2023;10: 1151028. doi:10.3389/fcvm.2023.1151028

100. Lelovas PP, Kostomitsopoulos NG, Xanthos TT. A Comparative Anatomic and Physiologic Overview of the Porcine Heart. J Am Assoc Lab Anim Sci. 2014;53: 432–438.

101. Crick SJ, Sheppard MN, Ho SY, Gebstein L, Anderson RH. Anatomy of the pig heart: comparisons with normal human cardiac structure. J Anat. 1998;193: 105–119. doi:10.1046/j.1469-7580.1998.19310105.x

102. Clark-Greuel JN, Connolly JM, Sorichillo E, Narula NR, Rapoport HS, Mohler ER, et al. Transforming Growth Factor-β1 Mechanisms in Aortic Valve Calcification: Increased Alkaline Phosphatase and Related Events. The Annals of Thoracic Surgery. 2007;83: 946–953. doi:10.1016/j.athoracsur.2006.10.026

