## Supplemental Figures for "Trymethylamine-N-oxide, a gut-derived metabolite, induces myofibroblastic activation of valvular interstitial cells through endoplasmic reticulum stress"

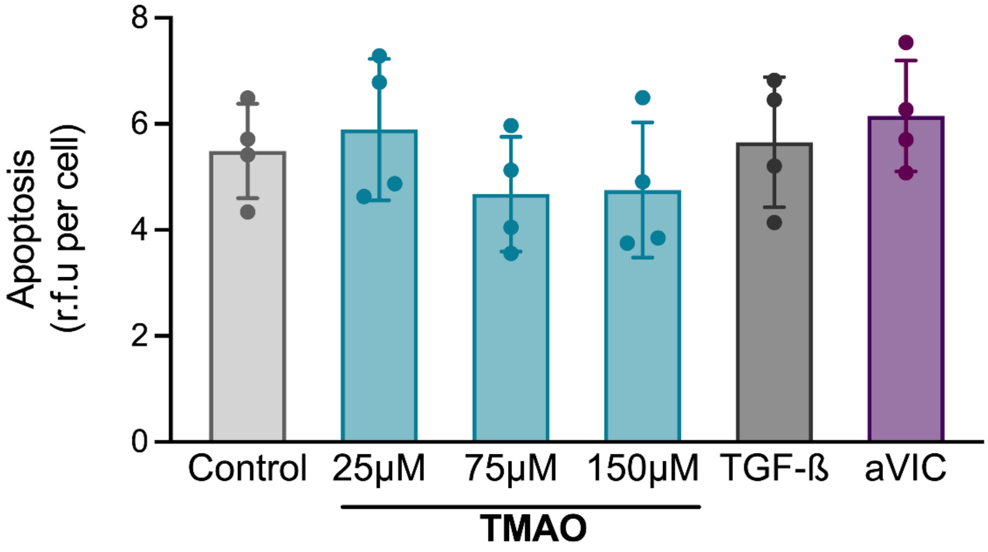


**Supplementary Figure 1. TMAO has no effect on qVIC apoptosis.** Evaluation of qVIC apoptosis levels after 3 days of TMAO treatment through a caspase 3/7 activity assay. No statistically significant differences were identified between experimental groups.

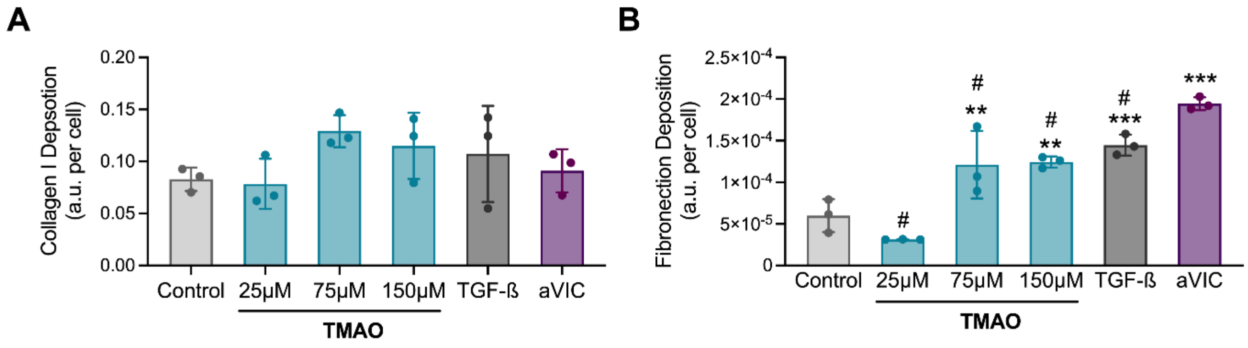


**Supplementary Figure 2. Effects of TMAO on ECM protein deposition: collagen and fibronectin.** Quantification of (A) collagen and (B) fibronectin deposition by qVICs treated with TMAO for 3 days via *in situ* ELISAs. n = 3-5 replicates per condition. **$p$<0.005, ***$p$<0.001 compared with the control, #$p$<0.05 compared to aVICs.

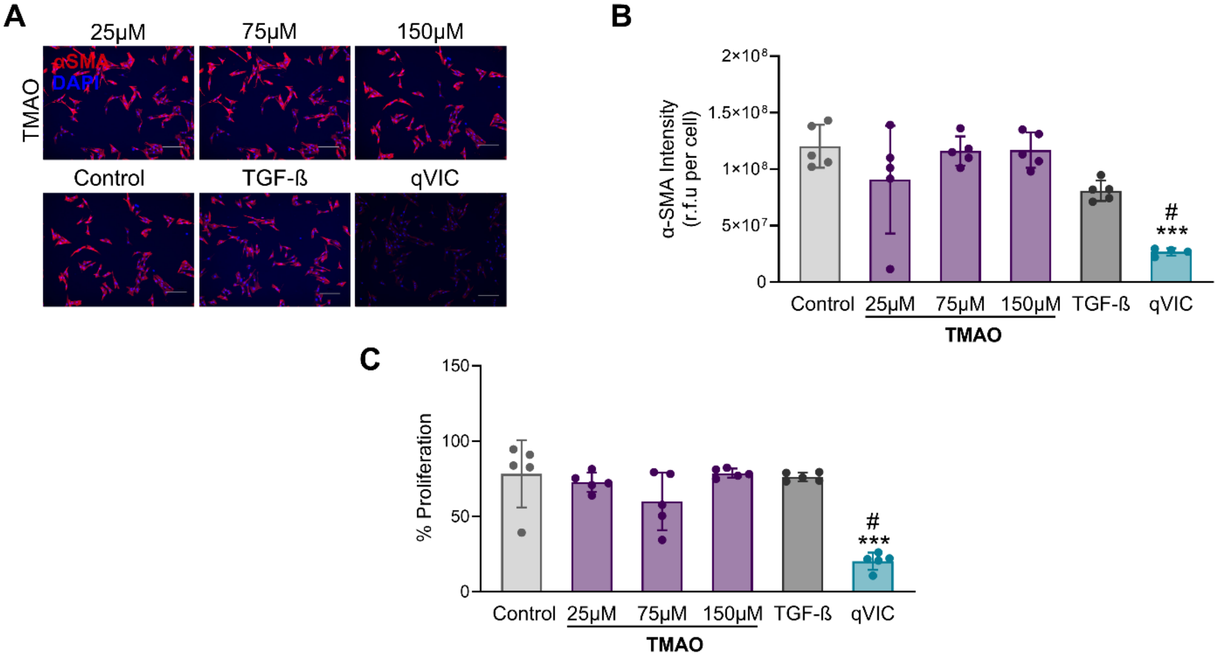


**Supplementary Figure 3 aVICs are not sensitive to low concentrations of TMAO.** Conventionally cultured aVIC were treated with TMAO (25-150μM) for 3 days. (A) Immunocytochemistry staining for αSMA (red). Cell nuclei are stained blue. (B) Quantification of αSMA staining intensity in (A). (C) Quantification of the percentage of EdU-positive proliferating cells. Scale bars represent 200 μm. n = 3-5 replicates per condition. One-way ANOVA followed by Tukey's multiple comparisons test. **p<0.005, ***p*<0.005, ****p*<0.001 compared with the control. #*p*<0.01 compared to treatment with TGF-$ß_1$.

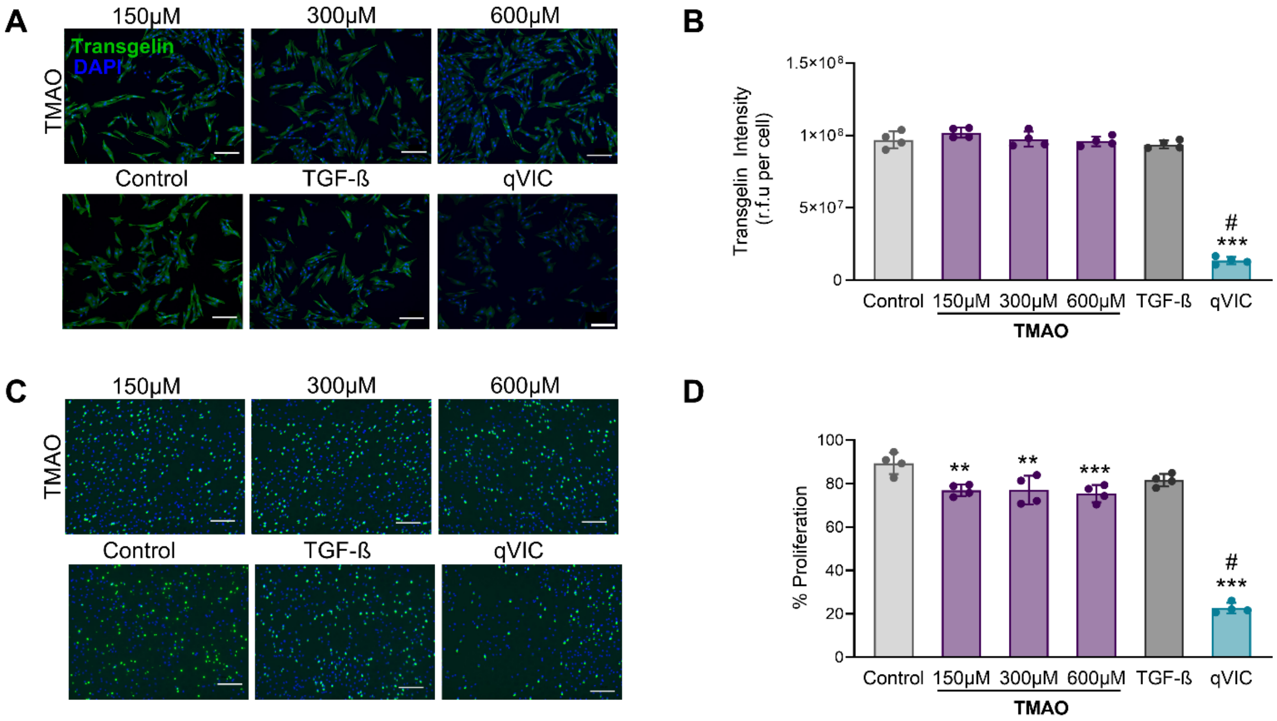


**Supplementary Figure 4. Effects of TMAO on transgelin expression and proliferation in conventionally cultured aVICs.** Conventionally cultured aVICs were treated with TMAO (150-600 μM) for 3 days. (A) Representative images of immunocytochemistry staining for transgelin (green). Cell nuclei are stained in blue. (B) Quantification of transgelin staining intensity in (A). (C) EdU staining to identify proliferating cells after 8 hours of incubation with EdU. (D) Quantification of the percentage of EdU-positive proliferating cells in (C). Scale bars represent 200 μm. n = 3-5 replicates per condition. One-way ANOVA followed by Tukey's multiple comparisons test. $^{**}p<0.005$, $^{***}p<0.001$ compared with the control. #$p<0.01$ compared to treatment with TGF-$ß_1$.

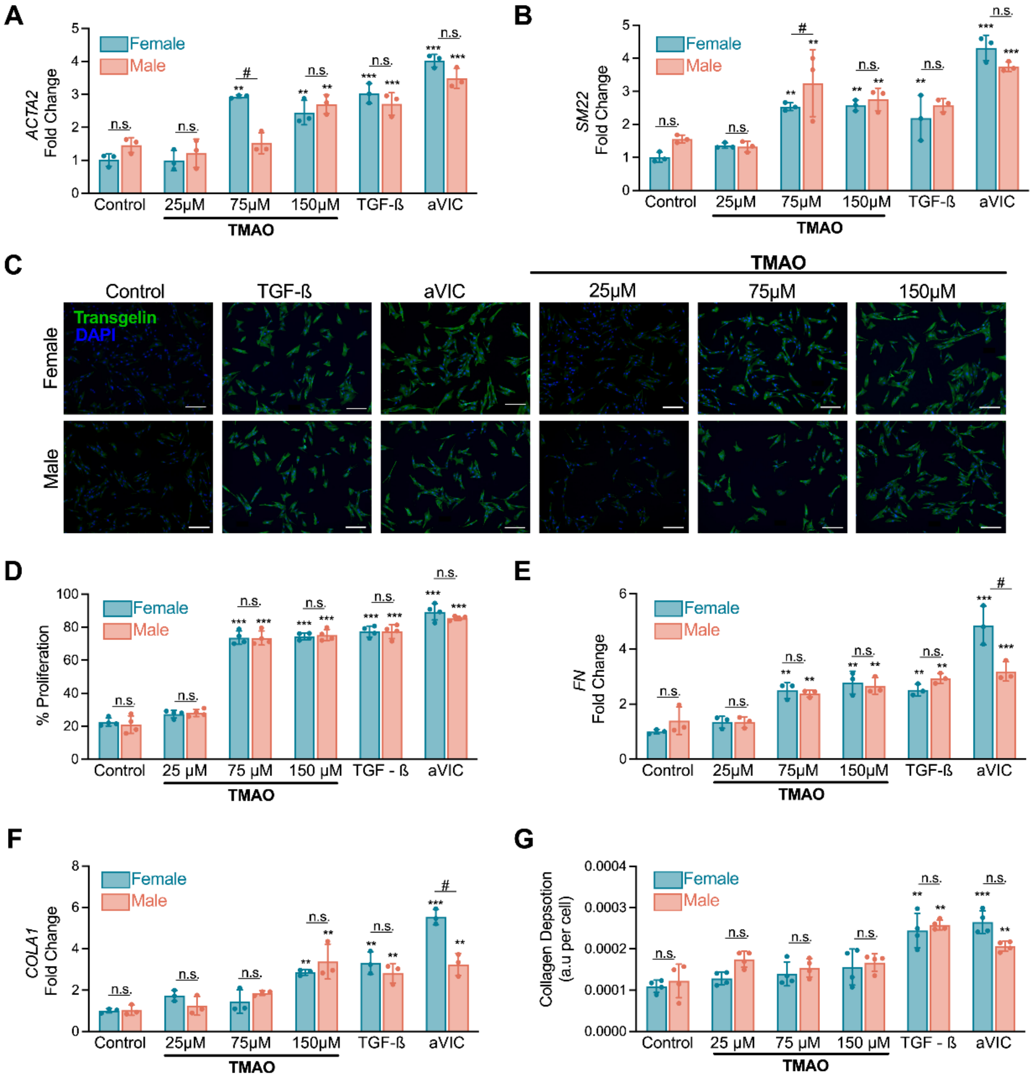


**Supplementary Figure 5. Sex-independent effects of TMAO on myofibroblastic activation and ECM protein production in qVICs.** Female and male qVICs were generated and treated with TMAO (25-150 μM) for 3 days. (A-B) Quantification of myofibroblastic gene expression levels for (A) *ACTA2* and (B) *SM22* via qRT-PCR. (C) Representative images of immunocytochemistry staining for transgelin (green). Cell nuclei are stained in blue. (D) Quantification of transgelin staining intensity in (C). (D) Quantification of the percentage of EdU-positive proliferating cells. (E-F) Gene expression analysis of (E) *FN* and (F) *COL1A1.* (G) Analysis of collagen deposition via *in situ* ELISA. Scale bars represent 200 μm. n = 3-5 replicates per condition. Two-way ANOVA was performed, followed by Tukey's multiple comparisons test. **$p < 0.005$, *$p < 0.001$ compared to the untreated control for the corresponding sex. n.s. denotes no statistically significant difference between male and female VICs for that condition. #$p<0.01$ for comparison shown.

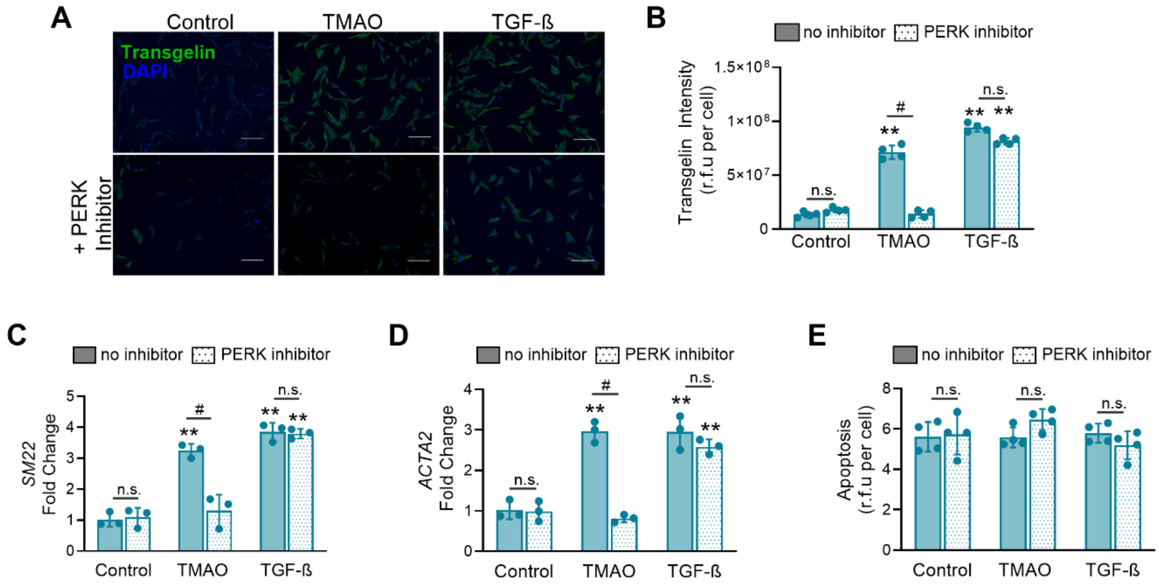


**Supplementary Figure 6. Effects of PERK inhibition on TMAO-induced qVIC activation and apoptosis in qVICs.** Female qVICs were pre-treated with GSK2656157, a PERK inhibitor, for 1-hour prior to treatment with TMAO. (A) Representative images of immunocytochemistry staining for transgelin (green). Cell nuclei are stained in blue. (B) Quantification of transgelin staining intensity in (A). (C-D) Quantification of myofibroblastic gene expression levels for (C) *SM22* and (D) *ACTA2.* (E) Evaluation of apoptosis levels through a caspase 3/7 activity assay. . Scale bars represent 200 µm. n = 3-5 replicates per condition. Two-way ANOVA, followed by Tukey's multiple comparisons test. $^{**}p<0.005$, $^{***}p<0.001$ compared with the control. n.s. denotes no statistically significant differences found between samples treated with and without the PERK inhibitor. #$p<0.001$ indicates a significant difference between samples treated with the PERK inhibitor and those without.

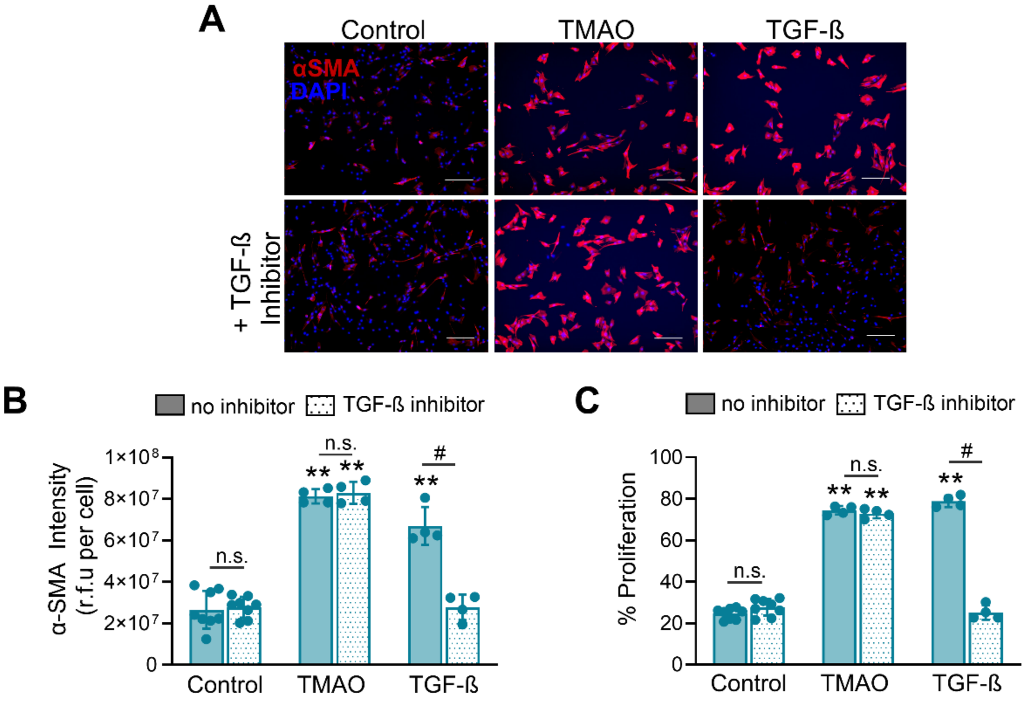


**Supplementary Figure 7. Inhibition of the TGF-ß$_1$ receptor does not impact TMAO-induced activation.** Female qVICs were pre-treated with SB431542, a TGF-ß$_1$ inhibitor that blocks activin receptor-like kinases on the cell surface, for 1 hour before treatment with TMAO. (A) Immunocytochemistry staining for αSMA (red). Cell nuclei are stained blue. (B) Quantification of αSMA staining intensity in (A). (C) Quantification of the percentage of EdU-positive proliferating cells. The scale bar represents 200 µm. n = 3-5 replicates per condition. Two-way ANOVA, followed by Tukey's multiple comparisons test. **p<0.005, ***p<0.001 compared with the control. n.s. denotes no statistically significant differences found between samples treated with and without the TGF-ß$_1$ inhibitor. #p<0.001 indicates a significant difference between samples treated with the TGF-ß$_1$ inhibitor and those without.
